# Small molecule enhancers of endosome-to-cytosol import augment anti-tumour immunity

**DOI:** 10.1101/2020.05.05.079491

**Authors:** Patrycja Kozik, Marine Gros, Daniel N. Itzhak, Leonel Joannas, Joao G. Magalhaes, Andrés Alloatti, Elaine Del Nery, Georg H.H. Borner, Sebastian Amigorena

## Abstract

Efficient cross-presentation of antigens by dendritic cells (DCs) is critical for initiation of anti-tumour immune responses. Yet, several steps of antigen intracellular traffic during cross-presentation are incompletely understood: in particular, the molecular mechanisms and the relative importance of antigen import from endocytic compartments into the cytosol. Here, we asked whether antigen import into the cytosol is rate-limiting for cross-presentation and anti-tumour immunity. By screening 700 FDA-approved drugs, we identified 37 import enhancers. We focused on prazosin and tamoxifen, and generated proteomic organellar maps of drug-treated DCs, covering the subcellular localisations of over 2000 proteins. By combining organellar mapping, quantitative proteomics, microscopy, and bioinformatics, we conclude that import enhancers undergo lysosomal trapping leading to membrane permeation and antigen release into the cytosol. Enhancing antigen import facilitates cross-presentation of both soluble and cell-associated antigens. Systemic administration of prazosin also led to reduced growth of MC38 tumours and to a synergistic effect with checkpoint immunotherapy in a melanoma model. Thus, inefficient antigen import into the cytosol limits antigen cross-presentation, restraining the potency of anti-tumour immune responses and efficacy of checkpoint blockers.

**Highlights:**

- Quinazolamine derivatives are potent enhancers of antigen import into the cytosol
- Antigen import enhancement occurs as a consequence of lysosomal trapping
- Antigen import enhancers facilitate cross-presentation and synergise with PD-L1
- Proteomic organellar maps of cDC1s as a tool to detect biological effects of drugs

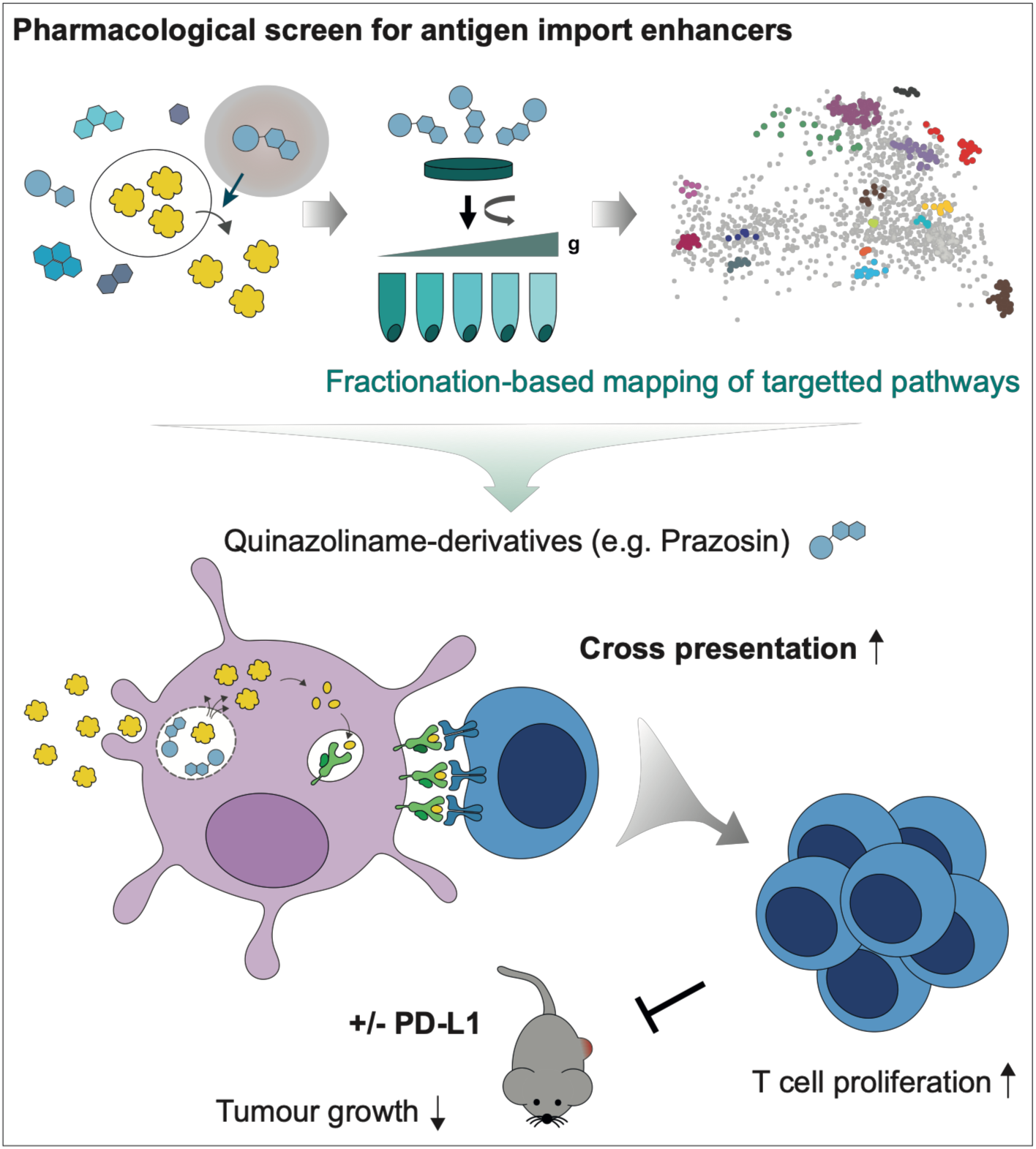

## Introduction

Accumulation of mutations in cancer is a key factor during disease progression, yet, it can also render cancer cells vulnerable to cytotoxic T cells (CTLs). T cell-mediated anti-tumour immune responses are primarily initiated by type 1 conventional dendritic cells (cDC1s) (Böttcher and Reis e Sousa, 2018). While these immune responses can in principle prevent or restrict tumour growth, they are usually not nearly as potent as responses against pathogens. In recent years, checkpoint inhibitors emerged as a promising tool to enhance anti-tumour immunity and were effective in providing long lasting remissions. Nevertheless, their efficacy is largely dependent on pre-exisiting immunity and the benefits are only seen in a fraction of patients (Crittenden et al., 2018). Therefore, a better understanding of the mechanisms and rate-limiting steps involved in priming of naive tumour-specific T cells will be critical for improving immunotherapeutic strategies.

Efficient priming relies on the delivery of three signals to naive T cells: signal 1 - relevant antigen (e.g. a mutated peptide) presented in the context of MHC class I; signal 2 - co-stimulatory molecules expressed by antigen presenting cells (APCs); and signal 3 - cytokines, which ultimately determine whether the response will lead to immunity or tolerance. Many approaches have been explored to deliver appropriate signals 2 and 3, including stimulating DCs maturation with a variety of TLR ligands (e.g. Poly(I:C) or CpG) or growth factors (e.g. FLT3L) (Brunner et al., 2000; Hammerich et al., 2019; Salmon et al., 2016; Sánchez-Paulete et al., 2018). However, increasing the efficiency of presentation of tumour antigens on MHC class I has proven more challenging.

Tumour antigens are presented by APCs via a process termed cross-presentation. Cross-presentation involves endocytic uptake of exogenous proteins followed by generation of short peptides that can be loaded onto MHC class I. Two models have been proposed to describe where peptide generation takes place (reviewed in (Gros and Amigorena, 2019)). In the vacuolar model, peptides are generated by endolysosomal proteases (primarily cathepsins) and directly loaded onto MHC class I (Shen et al., 2004). In the cytosolic model, antigens are imported into the cytosol, processed by the proteasome, and delivered into the lumen of MHC class I-containing compartments via the TAP transporter (Ackerman et al., 2003; Guermonprez et al., 2003; Kovacsovics-Bankowski and Rock, 1995). Considering differences in cleavage-specificities among the different proteases, the cytosolic model provides an attractive explanation of how APCs would generate peptides similar to those presented by target cells, where the majority of epitopes is also generated by proteasomes. Both TAP- and immunoproteasome-deficient mice are defective in cross-presentation (Palmowski et al., 2006; Rock and Shen, 2005), but whether these effects are indeed due to specific inhibition of cross-presentation, and whether the cytosolic pathway is dominant in vivo still requires verification. Similarly, mechanistic details of endosome-to-cytosol transport have remained elusive.

Irrespective of the precise mechanism, the importance of cross-presentation in initiation of anti-tumour responses has now been demonstrated in a variety of mouse models. cDC1s appear to be most efficient cross-presenters *in vivo* and Batf3-/-mice that lack cDC1s, do not mount efficient T cell responses (Hildner et al., 2008). In mice with a Wdfy4 deletion (Theisen et al., 2018) or a DC-specific knockout of Sec22b (Alloatti et al., 2017), cDC1s are present but deficient in the ability to cross-present. Both models are unable to prime naive T cells against tumour-associated antigens and fail to control tumour growth. Similar to cDC1-deficient mice (Sánchez-Paulete et al., 2016), Sec22b knockouts are also resistant to treatment with checkpoint inhibitors. These data argue for an important role of cross-presentation in anti-tumour immunity. Indeed, delivering tumour antigens to cross-presenting cells (e.g. via antibody-antigen conjugates), has been effective in promoting CTL responses (Bonifaz et al., 2002; Caminschi et al., 2008; Sancho et al., 2008). In the clinic, vaccination with long peptides comprising neoepitopes has also been successfully used to boost generation of tumour-specific T cells (Ott et al., 2017). These approaches of boosting antigen presentation are, however, costly to implement as they require prior identification of cancer neoantigens (e.g. through next generation sequencing of tumour samples).

Here, we present a strategy for enhancing efficiency of T cell priming, by facilitating antigen presentation by DCs. Our study was based on the hypothesis that import of internalised antigens into the cytosol might be limiting for the efficiency of cross-presentation. With this in mind, we set up an assay to screen a library of over 700 FDA-approved compounds to identify enhancers of antigen import. We demonstrated that these molecules indeed facilitated cross-presentation of both soluble and cell-associated antigens. To evaluate the biological activity of two import enhancers, prazosin and tamoxifen, we generated comprehensive proteomics-based organellar maps from treated and untreated cells. We established that our most potent compound, prazosin, has a highly specific effect on endolysosomal membrane permeability. This encouraged us to pursue *in vivo* studies, where we demonstrated that systemic administration of prazosin leads to better control of tumour growth and synergises with checkpoint-based anti-tumour immunotherapy.

## Results

### Selected ERAD inhibitors enhance antigen import

ER-associated protein degradation (ERAD) machinery has been proposed to play a key role in import of antigens from endosomes and phagosomes into the cytosol (Giodini and Cresswell, 2008; Imai et al., 2005; Zehner et al., 2015). Recently, however, we demonstrated that mycolactone, a potent inhibitor of Sec61 (a candidate ERAD translocon), does not inhibit antigen import (Grotzke et al., 2017). Here, we initially employed a pharmacological approach to evaluate the contribution of other ERAD components to antigen import. We selected a range of ERAD inhibitors and tested them using a β-lactamase-based antigen import assay (Figure 1A) (modified from (Cebrian et al., 2011). As a model system, we used the cell line MutuDC, which phenotypically corresponds to splenic cDC1s (Fuertes Marraco et al., 2012) (see also Figure 1G). To prevent tested compounds from affecting antigen uptake, we pulsed MutuDCs with β-lactamase for 3 hours, and subsequently treated them with the different inhibitors for 2 hours. To detect β-lactamase translocation into the cytosol, we loaded the cells with a cytosolic β-lactamase substrate, CCF4. When β-lactamase enters the cytosol, it cleaves the β-lactam ring in the CCF4 and disrupts FRET between its two subunits causing a shift in fluorescence from blue to green (Figure 1A). We monitored this change in fluorescence by flow cytometry (Figure 1B). The two compounds that target the ubiquitin pathway, PR-619 and Eeyarestatin I, inhibited antigen import (Figure 1B, consistent with previous data (Grotzke et al., 2017; Zehner et al., 2015). Unexpectedly, a p97 inhibitor, DbeQ, and a β-importin inhibitor, importazole, enhanced antigen import (Figure 1B, S1). This effect was not recapitulated with a more potent p97 inhibitor, NMS-873, suggesting it might be due to off-target activity. Hence, while these data highlight the role of the ubiquitin system in antigen import, they did not provide evidence supporting the role of other ERAD components. The dramatic enhancement of antigen import observed with two of the compounds suggests that antigen import is relatively inefficient, and that it maybe rate-limiting for cross-presentation.

**Figure 1.**
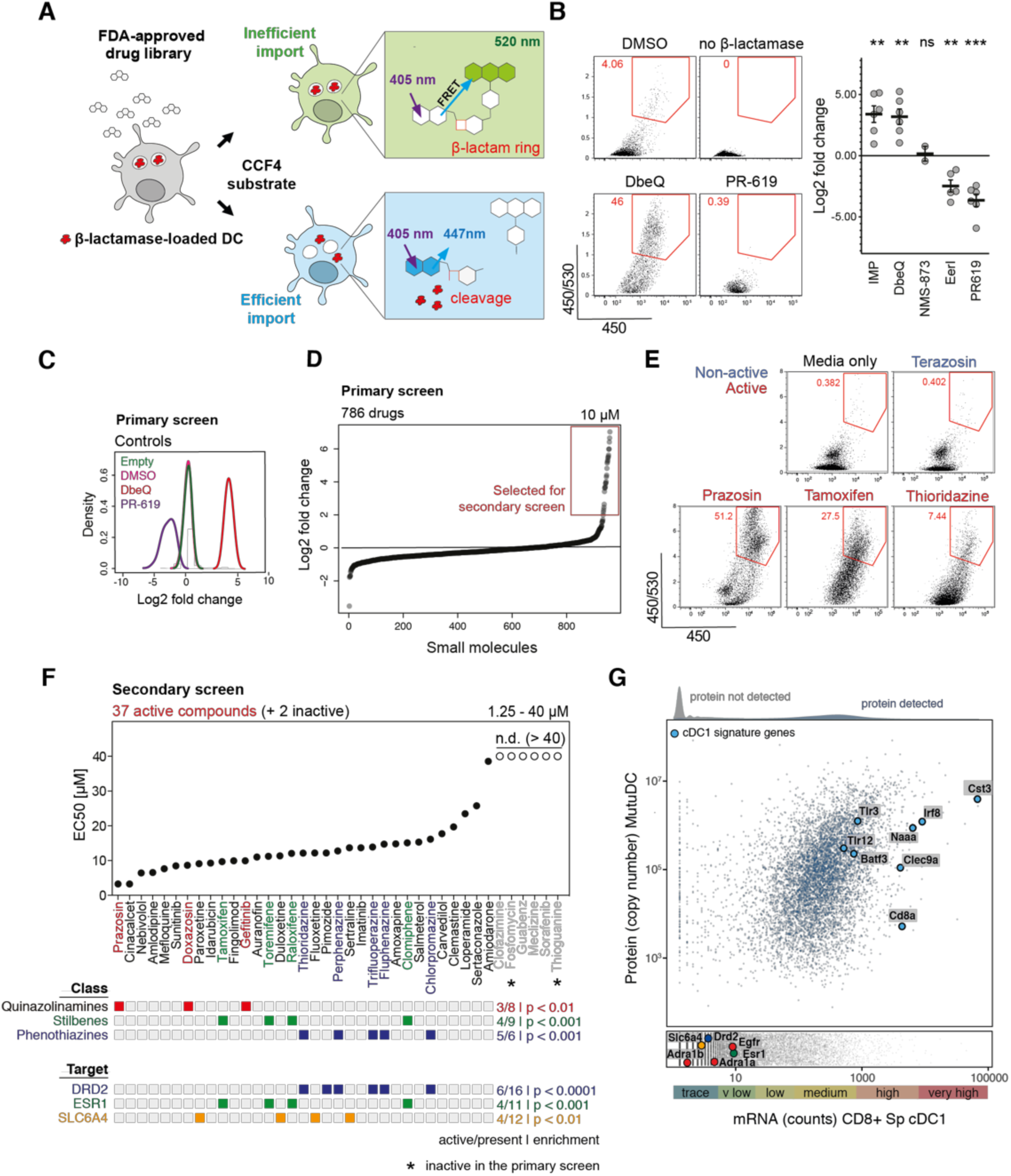
Small molecule screen to identify enhancers of antigen import into the cytosol. (A) Schematic representation of the β-lactamase assay used to monitor the efficiency of antigen import into the cytosol. MutuDCs were fed with β-lactamase for 3h followed by 2h incubation with small molecules (at 37°C). CCF4 loading was performed at RT for 1h, and followed by 16h incubation at RT to increase the sensitivity of the assay (Zlokarnik et al., 1998). Change in CCF4 fluorescence was monitored by flow cytometry. (B) Differential effects of ERAD inhibitors on antigen import into the cytosol. Representative flow cytometry data for selected ERAD inhibitors and quantification of the fold change in antigen import relative to DMSO controls. Within each condition, dots represents data from independent experiments, and lines represent means +/-SE. (C) Quality control of the FDA library screen. fold changes in the efficiency of antigen import (relative to DMSO) were calculated. The histograms show distribution of fold changes for each control (all wells across the ten 96-well plates). (D) Results from the FDA library screen. Fold changes in β-lactamase import for the 786 tested drugs. The screen was performed once and 37 compounds were selected for the secondary screen (highlighted with the red box). (E) Examples of the flow cytometry profiles in the antigen import assay for selected active and non-active compounds. (F) Results from the secondary screen of 37 compounds (and two control compounds, not active in the primary screen). Each drug was tested at five concentrations in two independent experiments. EC50 values (concentration required for 50% of maximal activity) values were calculates as described in Figure S2. Information about chemical classes and candidate targets was obtained from DrugBank database. The classes and targets enriched in the group of active vs non-active compounds are represented with coloured squares. The enrichment of targets for hits (compared to the entire library) was calculated using Fisher’s test. (G) Analysis of gene and protein expression in CD8^+^ cDC1s. mRNA expression data (RNAseq) for CD8^+^ splenic DCs was downloaded from the Immgen.org database and whole cell proteomic abundance data were generated by mass spectrometry from MutuDCs. 7427 proteins were detected and full absolute copy number proteome data of MutuDCs are available via the web resource (http://dc-biology.mrc-lmb.cam.ac.uk). The blue line indicates the threshold for expression. The points between the grey vertical lines represent proteins for which corresponding microarray probes were not identified in the Immgen dataset. The points between the grey horizontal lines represent genes for which the corresponding proteins were not detected. See also Figure S1, S2, Supplemental Data S1, S2.

### β-lactamase-based screen for enhancers of antigen import

Enhancement of antigen import by DbeQ and importazole established a proof of concept that this process can be pharmacologically manipulated, and prompted us to develop a screen for small molecule import enhancers. We performed the screen in MutuDCs using the β-lactamase-based antigen import assay, and treating the cells with the 786 small molecules from a library of FDA-approved drugs (Supplemental Data S1). DbeQ and PR-619 were used as controls on each plate to track data quality (Figure 1C). We selected 37 drugs that increased antigen import at least two-fold in the primary screen for follow up (Figure 1D, E). Two non-active compounds were also included as additional negative controls.). 32 out of these 37 compounds exhibited a dose-dependent effect in the secondary screen (4% hit rate) (Figure 1F and Figure S2); they included three classes of chemically related compounds: quinazolinamines (prazosin, doxazosin, and gefitinib), stilbenes (clomiphene, raloxifene, tamoxifen and toremifene), and phenothiazines (chlorpromazine, fluphenazine, perphenazine, thioridazine, trifluoperazine) (Figure 1F).

To understand the mechanism of antigen import enhancement, we first searched for common targets among the active compounds. Using the DrugBank database (Law et al., 2014), we identified previously described targets for 714 of the compounds present in the FDA library. Three of these targets were significantly enriched among active versus non-active compounds: estrogen receptor (Esr1), dopaminergic receptor (Drd2), and serotonin transporter (Slc6a4) (Figure 1F lower panel). However, none of these three proteins is actually expressed in CD8^+^ cDC1s according to Immgen.org mRNA expression data (Yoshida et al., 2019); they are also not present among the 7,427 proteins we detected in MutuDC by proteomics (Figure 1G, Supplemental Data S2). Considering that out of 11 estrogen receptor modulators present in the library, antigen enhancement was only observed for inhibitors from the stilbene family, the enrichment appeared to be linked to the structure of these compounds, rather than to the inhibition of known targets. Similarly, no protein and only trace mRNA were detected for targets of the most potent class of enhancers identified, quinazolinamines (Adra1, Adra2, and Egfr). Interestingly, both DbeQ and importazole belong to the same chemical family; hence, half of the ten quinazolinamine derivatives tested in this study facilitated import of internalised antigens, despite being marketed as inhibitors of different targets.

### Organellar maps to determine biological activity of small molecules in DCs

A variety of “hidden phenotypes” and promiscuous effects have been observed for numerous clinically approved drugs (MacDonald et al., 2006). These additional phenotypes can often be beneficial for novel therapeutic indications, yet there are few approaches to detect the cellular effects of a compound in an unbiased manner. To characterise the mechanism of antigen import enhancement, we developed a generic strategy to evaluate the biological activity of pharmacological compounds through comparative spatial proteomics (Figure 2A). Many if not most cell biological processes are accompanied by protein subcellular localisation changes (Lundberg and Borner, 2019). Hence, we adapted our previously developed method for generating organellar maps, to pinpoint the subcellular localisations of thousands of proteins in a single experiment (Itzhak et al., 2017; 2018). The comparison of organellar maps made under different physiological conditions allows the capture of drug induced protein translocations (Itzhak et al., 2016), and thus provides a universal and scalable tool for inferences about cellular responses and drug targets.

**Figure 2.**
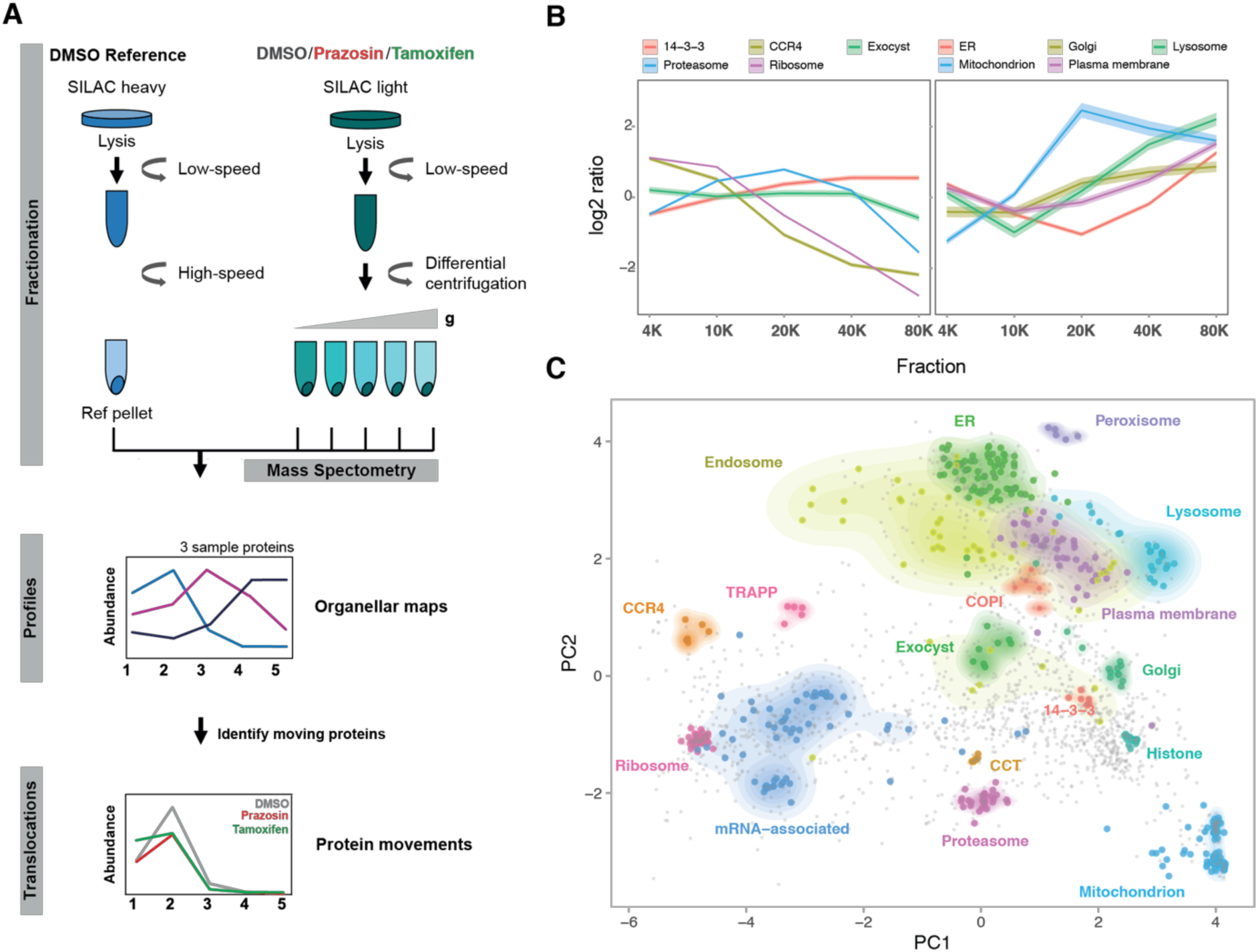
Organellar mapping in dendritic cells. (A) Schematic representation of the fractionation profiling approach for making organellar maps. Metabolically labelled (SILAC heavy - vehicle treated, and light - vehicle or drug treated) MutuDCs are lysed mechanically. Post-nuclear supernatant from light labelled cells is subjected to a series of differential centrifugation steps, to separate organelles partially. In parallel, post-nuclear supernatant from heavy labelled cells is pelleted at high speed to obtain a reference fraction, which is spiked into each of the light fractions. Quantitative mass spectrometry allows the accurate determination of abundance distribution profiles across the light subfractions for individual proteins. Proteins associated with the same organelle have similar profiles, and different organelles have distinct profiles. Principal Component Analysis is used to visualize organellar clusters. (B) Examples of the log2 heavy/light ratios for proteins in selected organelles and protein complexes from vehicle treated MutuDCs (mean +/- 95% CI). (C) Organellar maps of MutuDCs visualized by Principal Component Analysis. The first two principal components account for >90% of the variability in the data. Marker proteins of various organelles and known protein complexes are shown with coloured circles; density gradients for proteins in each cluster are also highlighted. See also Figure S3 and Supplemental Data S2.

To generate organellar maps, we separated post-nuclear supernatants from MutuDCs into five pellets obtained by differential centrifugation (Figure 2A). Each pellet was mixed 1:1 with a SILAC heavy ‘‘reference’’ membrane fraction and the samples were analysed by MS. To generate abundance profiles, we calculated heavy-to-light ratio for each protein in each fraction. Using organellar markers we previously established for HeLa cells (Itzhak et al., 2016), we confirmed clustering of proteins from different organelles (e.g. lysosome, peroxisome, and mitochondria) and protein complexes (e.g. proteasome, CCR4-NOT complex) (Figure 2B, 2C). These maps cover over 2000 proteins expressed in DCs and can be mined for protein subcellular localisation, absolute abundance (copy numbers and cellular concentrations), as well as nearest neighbours (i.e. potential interaction partners) via a web resource (http://dc-biology.mrc-lmb.cam.ac.uk, see also Figure S3, Supplemental Data S2).

We focused on two import enhancers from different chemical classes, prazosin (quinazolinamine) and tamoxifen (stilbene). To investigate their effects on organellar dynamics, we prepared maps from drug or vehicle-treated MutuDCs in biological duplicates (six maps in total; Supplemental Data S3). To detect significant protein translocations, maps of control and drug-treated cells were compared using MR (Movement and Reproducibility) plot analysis (Figure 3A). Tamoxifen treatment led to spatial rearrangements of 56 proteins in MutuDCs, whereas prazosin was affected only 33 proteins. The majority of prazosin hits (27/33) mapped to the lysosomal compartment (Figure 3B). These hits comprised 23 out of 24 detected soluble lysosomal enzymes (e.g. cathepsins) as well as three transmembrane proteins. Out 13 proteins shifting with both drug treatments, 12 also mapped to lysosome (for other lysosomal proteins, the M scores in the tamoxifen sample were just below the threshold). Other proteins that shifted upon tamoxifen treatment included components of COPI vesicles, stress granules (e.g. Caprin1, G3bp1), or CCR4-NOT complex (Figure 3B, Supplemental Data S3). Thus, in dendritic cells, tamoxifen has pleiotropic effects and prazosin is highly specific, but there is a common effect of both compounds on lysosomal proteins.

**Figure 3.**
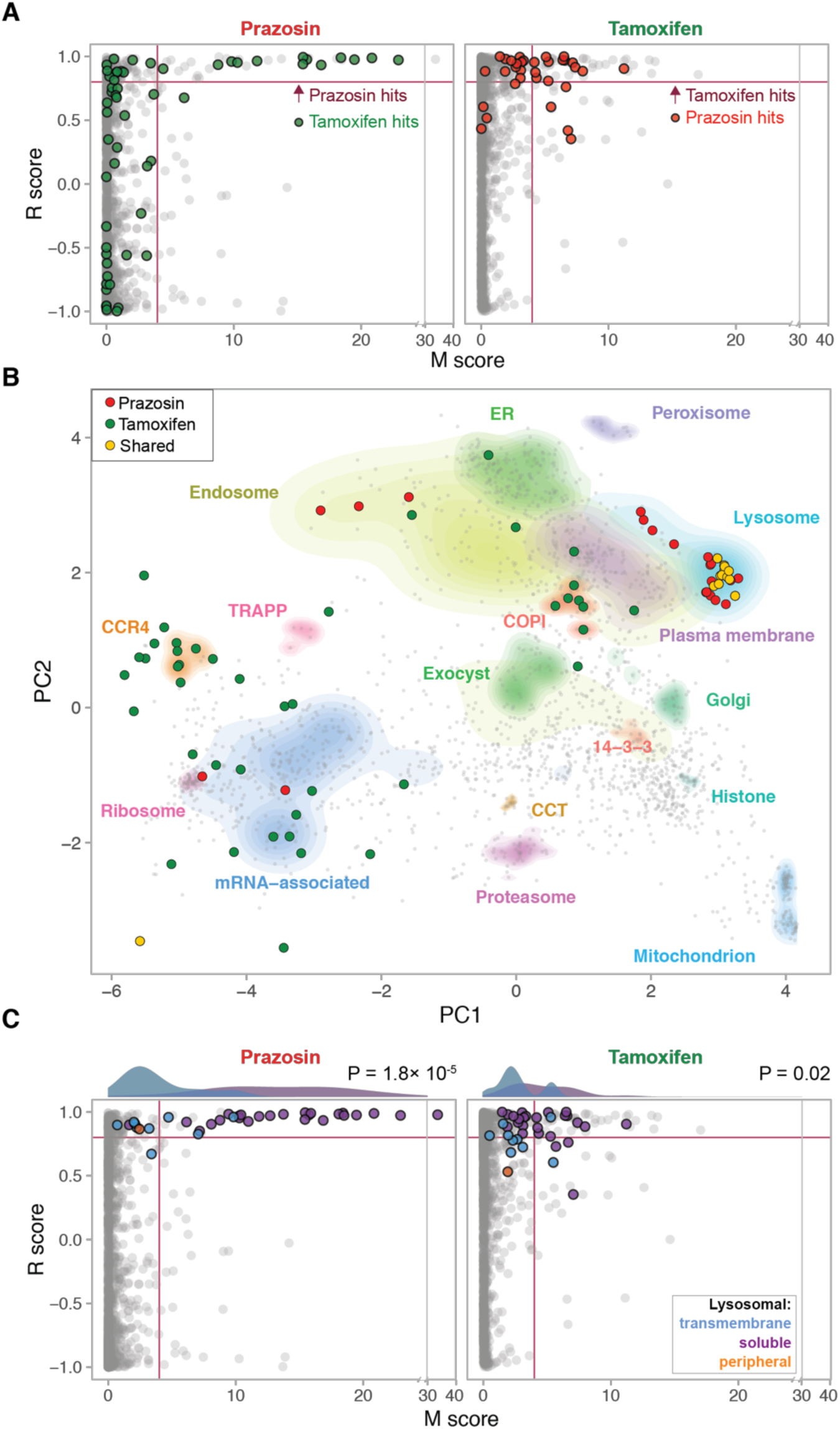
Dynamic organellar mapping to identify the subcellular changes in protein distribution upon drug treatment. MutuDCs were treated with either prazosin, tamoxifen or DMSO (control) for 4 h, in biological duplicate and samples were processed as described in Figure 2A. Statistical comparison of organellar maps made with different treatments was performed to identify proteins with profile shifts/altered subcellular localisation. (A) Drug-induced shifts in protein subcellular localization detected using a ‘MR’ plot analysis. For each protein, the movement (M score) and the reproducibility of the movement (R score) between maps was determined. Purple lines indicate cut-offs for significance. In prazosin plot the hits from tamoxifen treatment are shown for comparison (and vice-versa). (B) Shifting proteins from tamoxifen and prazosin-treated samples represented on the organellar map of MutuDCs. (C) MR plot highlighting all detected lysosomal proteins (soluble, transmembrane and peripheral (located on the cytosolic side of the membrane)). The histograms show distribution of the M scores for transmembrane and soluble lysosomal proteins. P values were calculated using the Mann-Whitney test.

We went on to analyse the overall behaviour of lysosomal proteins detected in MutuDCs in more detail. While the majority of soluble lysosomal proteins had high M scores (shift to the right of the MR plot), lysosomal transmembrane proteins show little or no translocation (Figure 3C). This difference in behaviour of soluble and transmembrane proteins suggests that lysosomal contents are either secreted into the extracellular space or leaked into the cytosol.

### Prazosin and tamoxifen induce lysosome permeability

To determine whether lysosomal contents in prazosin and tamoxifen treated cells are secreted or leaked, we performed quantitative proteomic analyses of whole cell extracts and cytosolic fractions (Supplemental Data S3). We observed significantly elevated levels of lysosomal enzymes in the cytosol of prazosin and tamoxifen-treated MutuDCs relative to control cells (Figure 4A). Since the levels of these proteins were not changed in whole cell proteome (Figure 4A, S4), we concluded that both prazosin and tamoxifen facilitate lysosomal leakage. Similar to what we observed using organellar maps, the prazosin effect is mostly restricted to lysosomal proteins, whereas tamoxifen affects a larger and more diverse set of proteins.

**Figure 4.**
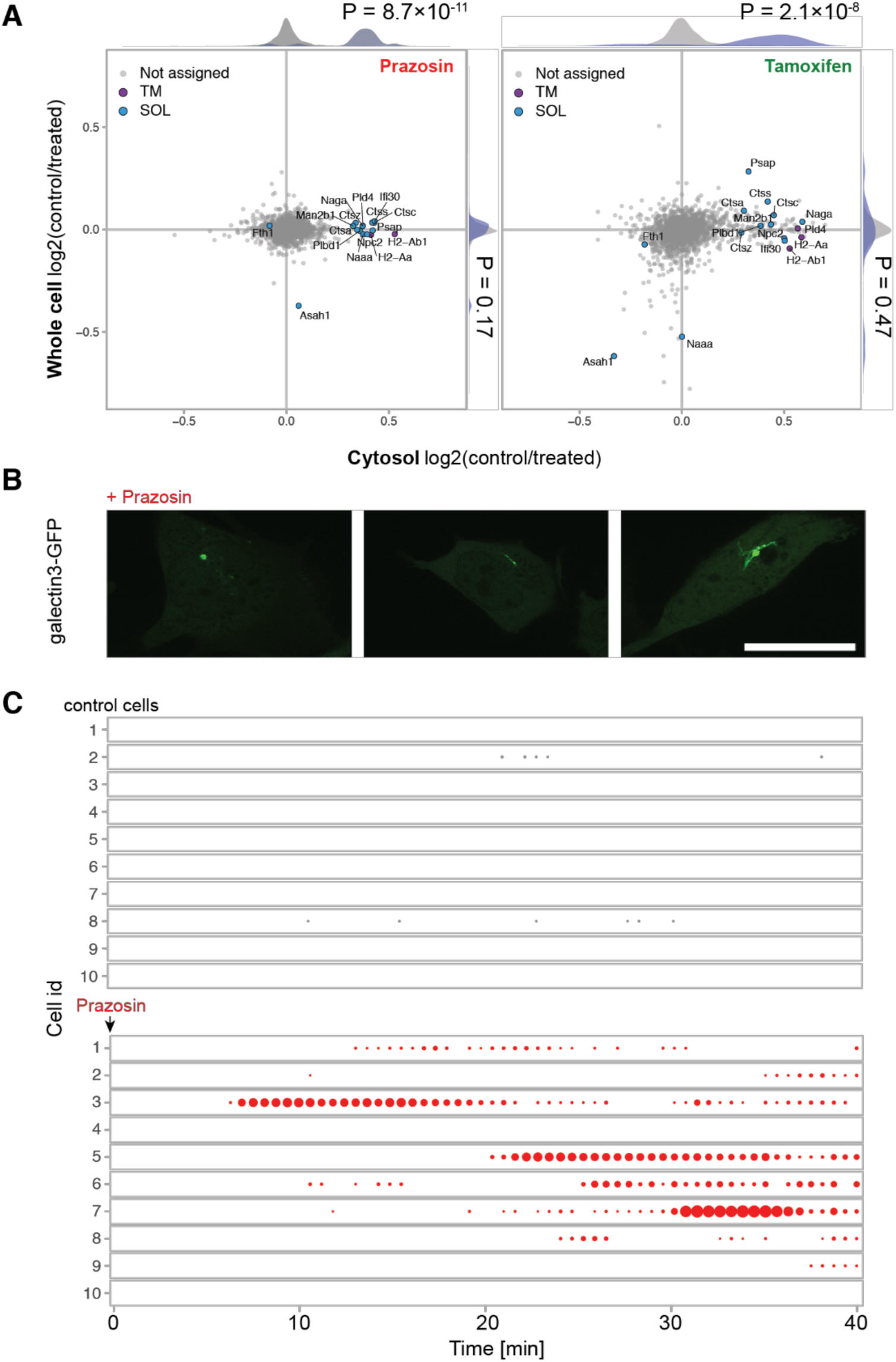
Prazosin and tamoxifen lead to lysosomal leakage. (A) Analysis of whole cell proteome and cytosol fractions from MutuDCs treated with prazosin, tamoxifen, or DMSO (control) for 4 h. The relative abundance of proteins from prazosin or tamoxifen vs vehicle-treated cells in whole cell vs cytosol proteomes. The histograms show distributions of all (grey) vs lysosomal (blue) proteins. P values were calculated using the Kolmogorov–Smirnov test. n=2 for whole cell lysates (SILAC quantification), n=4 for cytosol fractions (label-free quantification)). (B, C) MutuDCs stably expressing Galectin3-YFP were treated with prazosin and imaged continuously for 40 min. (B) Examples of Galectin3-YFP-positive structures observed in prazosin-treated cells. (C) Quantification of Galectin3-YFP recruitment in control and prazosin-treated cells. Circle sizes correspond to the area of Galectin3-YFP spot(s) in each cell at each time point. Representative data from ten cells imaged in one of two independent experiments. See also Figure S4 and Supplemental Movie S4.

To rule out that the observed lysosomal leakage was caused by increased compartment fragility and enhanced rupture during cell fractionation, we tested permeability of endolysosomal compartments *in vivo*. To this end, we used galectin-3-YFP probe and video microscopy. Galectin-3 is a cytosolic protein that associates with the carbohydrates on the luminal side of the endolysosomal compartments when membranes are damaged (Thurston et al., 2012). In control cells, galectin-3 signal is diffuse and galectin-3-positive structures are rarely observed (Supplemental Movie S4). Following addition of prazosin, however, there are frequent bursts of galectin-3-YFP recruitment to vesicular and tubular compartments in MutuDCs (Figure 4B, 4C, and Supplemental Movie S4). The morphology of these compartments is reminiscent to the endolysosomal structures previously observed in prazosin-treated cells (Zhang et al., 2012). The accessibility of the lysosomal lumen to a cytosolic probe demonstrates that membranes become permeable to proteins. In summary, we conclude that both prazosin and tamoxifen target lysosomes, causing membrane destabilisation and leakage of lysosomal contents, including internalised antigens, into the cytosol.

### Lysosomotropic properties of quinazolinamines mediate import enhancement

Considering that in dendritic cells, prazosin had a highly specific effect on lysosome permeability, we hypothesised that the enhancement of antigen import might be mediated through lysosomal trapping of the drugs. Lysosomal trapping occurs when a compound readily crosses membranes at neutral pH, but becomes protonated and membrane impermeable at acidic pH (Figure 5A, inset). This phenomenon has been observed for several classes of amine group-containing, amphiphilic compounds (Nadanaciva et al., 2011). Interestingly, all but one (auranofin) of the hits have physicochemical properties of lysosomotropic compounds, i.e. pKa between 6.5 and 11 and logP value greater than two (Figure 5A). We used BODIPY-conjugated prazosin to determine whether prazosin undergoes lysosomal trapping (Figure S5). Indeed, within seconds following addition, prazosin-BODIPY rapidly accumulated in vacuolar compartments in MutuDCs (Figure 5B). To test whether lysosomal trapping is required for antigen import enhancement, we performed the β-lactamase import assay in the presence of NH_4_Cl to neutralise the lysosomal pH. For all four compounds tested (prazosin, tamoxifen, DbeQ, and importazole), the enhancement of β-lactamase import was completely abolished in the presence of NH_4_Cl (Figure 5C). Therefore, the four drugs require low endocytic pH to enhance antigen import.

**Figure 5.**
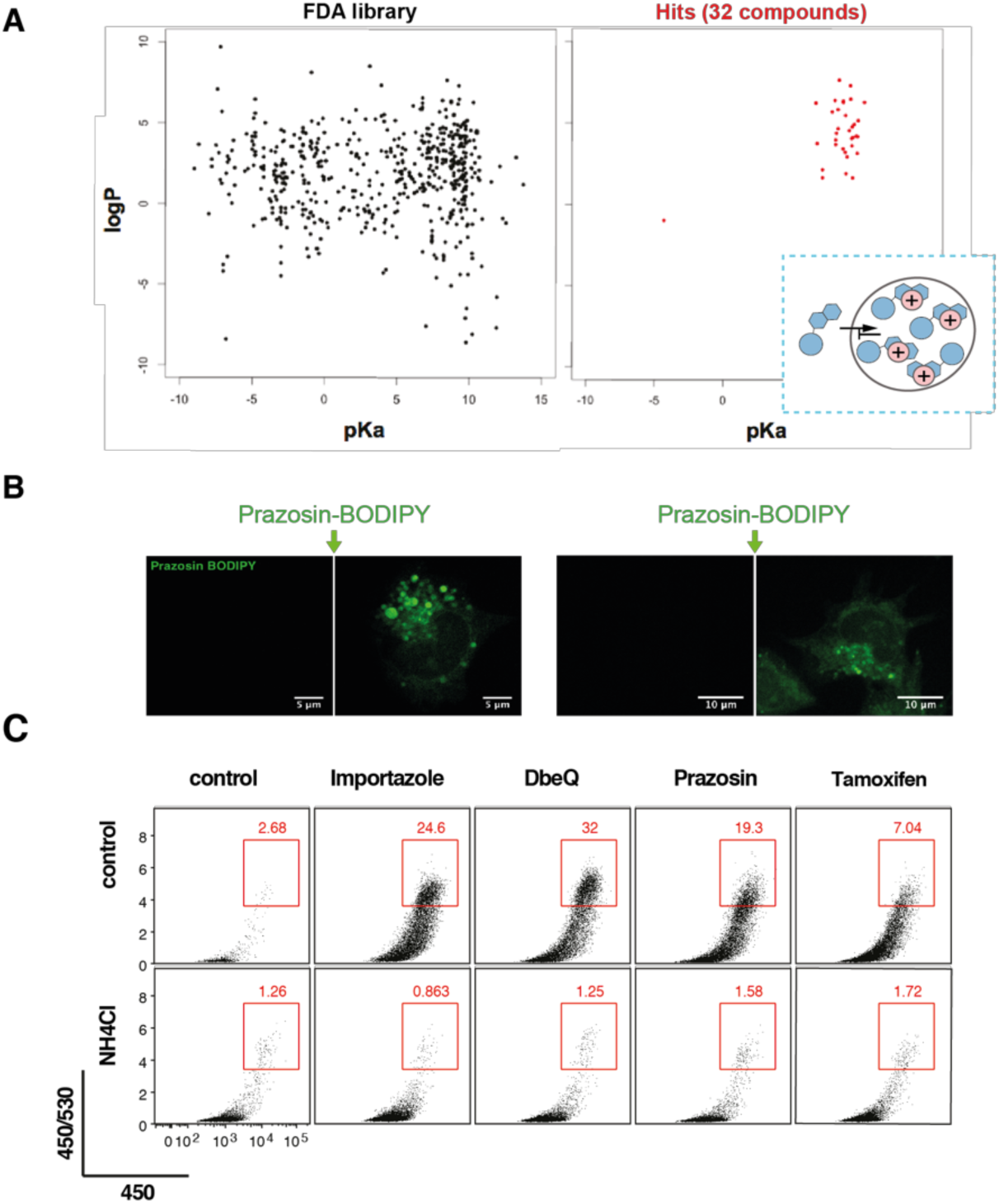
Antigen import enhancement is a consequence of lysosomal trapping. (A) Physicochemical properties (according to DrugBank data) of all the compounds present in the FDA library and of those active in the antigen import assay (yEC50 < 40 μM, see Figure 2d). All but one of the hits have physicochemical properties similar to those of lysosomotropic compounds (blue box). In the inset, schematic representation of the mechanism of lysosomal trapping. Membrane permeable small molecules diffuse freely across the membrane of acidic compartments. Protonation of weakly basic residues decreases membrane permeability and leads to accumulation of protonated compounds in the endo-lysosomal lumen. (B) MutuDCs cells were imaged immediately before and after addition of 5 μM prazosin-BODIPY. Data representative from three independent experiments. (C) Antigen import assay (Figure 2A) was performed in the presence of prazosin, importazole, tamoxifen and DbeQ with or without NH_4_Cl. Representative plots are shown (prazosin, n = 3; tamoxifen and DbeQ, n = 2) See also Figure S5.

### Antigen import enhancers augment cross-presentation and cross-priming

To determine if enhanced antigen import results in increased antigen cross-presentation to CD8^+^ T cells, we fed DCs with soluble ovalbumin (sOVA) before treatment with the drugs and incubation with K^b^/OVA_257-264_-specific B3Z T cell hybridoma. Both prazosin and tamoxifen treatment led to a dramatic, concentration-dependent enhancement of B3Z activation (Figure 6A). We observed a similar enhancement of cross-presentation of cell associated antigens (Figure 6B); in these experiments we co-cultured 3T3s (K^d^) expressing cytosolic OVA with MutuDCs for 5h and fixed the co-cultures before addition of the B3Z hybridomas. As demonstrated using a membrane labelling dye (PKH-26^+^), prazosin did not increase uptake of cell-associated material (Figure 6C). Importantly, we were also able to enhance cross-presentation of endotoxin-free sOVA (Figure 6D), which is normally not cross-presented efficiently due to the absence of TLR ligands (Alloatti et al., 2016; Burgdorf et al., 2008). This enhancement was not due to prazosin-mediated DC activation as we did not observe up regulation of activation markers in prazosin-treated DCs (Figure 6E). Finally, in accordance with the proposed mechanism of antigen import enhancement, we did not observe an increase in cross-presentation efficiency in the presence of NH_4_Cl (Figure 6F). Importantly, prazosin did not enhance T cell proliferation when DCs were pulsed with the short peptide SIINFEKL, indicating that it does not affect the general ability of DCs to prime T cells (Figure 6F). Together, these data indicate that facilitating antigen import into the cytosol overcomes the requirement for DC activation during cross-presentation, and suggest that antigen import might be a key regulatory step that determines which antigens are destined for cross-presentation.

**Figure 6.**
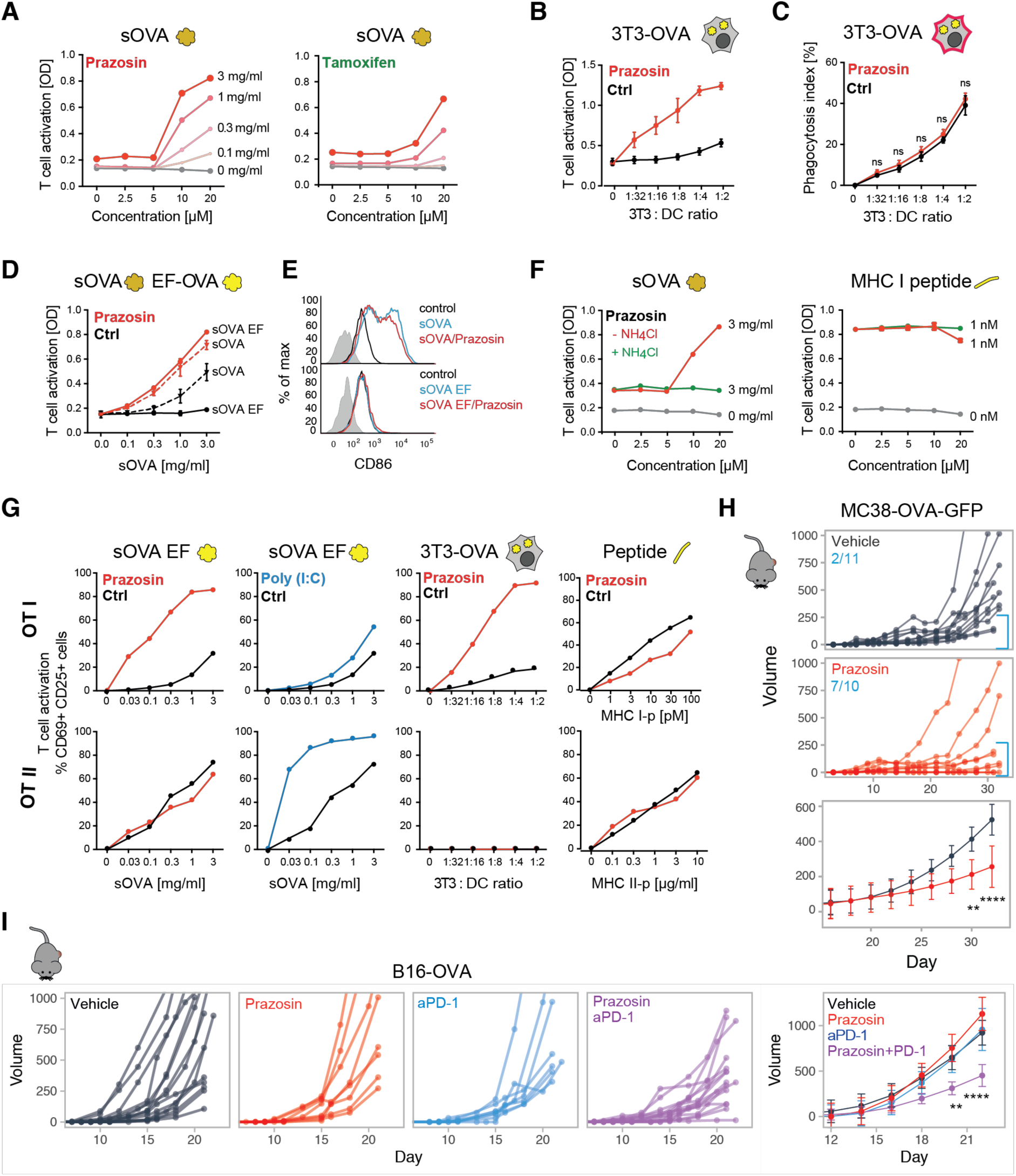
Prazosin enhances cross-presentation and cross-priming. (A) Antigen cross-presentation assay with B3Z hybridoma in the presence of increasing concentrations of prazosin or tamoxifen. The cells were pulsed with sOVA for 45 min, followed by 3.5 h incubation in the presence of the indicated compounds. Representative of three independent experiments. (B) The effect of prazosin on cross-presentation of cell-derived antigens. 3T3 cells expressing cytosolic OVA were used as antigen source and co-cultured with MutuDCs in for 5h in the presence or absence of prazosin. Mean from three independent experiments +/- SE. (C) Phagocytosis efficiency in the presence and absence of prazosin. 3T3s were labelled with PKH26, and acquisition of the dye by MutuDCs was analysed after 2h of co-culture. Mean from three independent experiments +/- SE. (D) MutuDCs were incubated with sOVA or sOVA EF for 45 min followed by 3.5 h incubation with prazosin. B3Z assay was used to monitor cross-presentation efficiency (representative plot from three independent experiments, error bars indicate SEM from technical duplicates). (E) For the analysis of DC activation, MutuDCs were incubated with sOVA/EF sOVA in the presence and in the absence of prazosin for 5 h, washed, further incubated for 16 h at 37°C, and stained with anti-CD86 (gated for live cells only). (F) Antigen cross-presentation assay with B3Z hybridoma in the presence of increasing concentrations of prazosin with or without 10 mM NH_4_Cl was performed as in (A). Mean +/- SE. (G) The effect of prazosin on antigen presentation to OT-I and OT-II cells. MutuDC were incubated with sOVA EF, OVA-expressing 3T3s or MHC class I or II peptides in the presence or absence of prazosin or Poly(I:C). (H) Tumour growth. Mice were injected s.c. with the MC38-OVA tumour cell. When tumours became detectable, the animals were treated systemically (IP) with 0.5 mg prazosin or vehicle control, 3 x week. Mice pooled from two independent experiments. The numbers indicate number of mice with tumours smaller than 250 mm3 at the end of the experiment. Lower panel represents best-fit curves for control and prazosin-treated groups, where means and SD were calculated using loess regression, the statistical significance was calculated using ANOVA with FDR Benjamini-Hochberg correction. ** p < 0.01 ****, p < 0.0001 (I) Tumour growth curves for mice injected s.c. with the B16-OVA tumour cells. From the day when tumours became detectable, mice were treated three times per week with 0.5 mg prazosin, 150 μg anti-PD-1, or the combination of both. Mice pooled from three independent experiments. The last panel represents best-fit curves for all groups, where means and SD were calculated using loess regression. The statistical significance was calculated using ANOVA with FDR Benjamini-Hochberg correction. ** p < 0.01, **** p < 0.0001.

We went on to determine if endosomal processing of antigens for presentation on MHC class II was also enhanced by prazosin. We fed DCs with soluble endotoxin-free OVA, cell-associated OVA, or the appropriate peptides that bind directly to MHC molecules and co-cultured them with OT-I (CD8^+^) and OT-II (CD4^+^) T cells (specific for K^b^/OVA_257-264_ and I- A^b^/OVA_323-339_ respectively). As shown in Figure 6G, presentation of soluble and cell-associated antigens to CD4^+^ OT-II cells was not affected by prazosin. This was in clear contrast with Poly(I:C), which strongly enhanced sOVA presentation to CD4^+^ but not to CD8^+^ T cells. Again, prazosin did not enhance T cell priming when DCs were treated with the short peptides directly, which would not require import into the cytosol for presentation. In summary, prazosin enhances antigen cross-presentation selectively and independently of DC maturation.

Finally, we investigated whether prazosin could be used to enhance antigen cross-presentation and anti-tumour immunity *in vivo*. In mice bearing MC38-GFP-OVA tumours, we observed reduced tumour growth following systemic treatment with prazosin (Figure 6H). In the more aggressive tumour model B16-OVA, prazosin alone was insufficient to control tumour growth and neither was a checkpoint inhibitor, anti-PD-1. However, combination of prazosin and anti-PD-1 led to a synergistic effect and a significant delay in tumour growth (Figure 6I). We conclude that a combination of checkpoint blockade and increased antigen cross-presentation can overcome resistance of certain tumours to immunotherapy.

## Discussion

In this study, we developed a strategy to harness the natural capacity of DCs to cross-present antigens by modulating a specific step involved in antigen processing: import into the cytosol. To enhance antigen import, we used small molecules identified through a pharmacological screen. We demonstrated that import is a rate-limiting step for cross-presentation, in particular for antigens free of pathogen-derived signals. This observation reinforces the hypothesis that import into the cytosol is a regulated step (Zehner et al., 2015), and implies that endogenous signals that drive import and cross-presentation in the absence of infection await identification.

Boosting antigen import and cross-presentation synergises with anti-PD-1-mediated immunotherapy in a tumour model unresponsive to the antibody alone, suggesting that cytosolic antigen cross-presentation plays a critical role in anti-tumour immune responses. Thus, enhancing antigen cross-presentation with small molecules provides a new strategy for combination therapies with checkpoint blockers. The major advantage of this approach, in comparison to tumour antigen-containing vaccines is that enhancing the natural capacity of DCs to route internalised antigens for cross-presentation does not require prior identification of specific epitopes.

Here we report over 30 FDA-approved small molecule enhancers of antigen import and we characterise the effects of two molecules (prazosin and tamoxifen) in more detail. Using a combination of proteomics, microscopy, and bioinformatics, we concluded that import enhancement occurs as a consequence of lysosomal trapping of the drugs, enhancing lysosomal permeability. A number of mechanisms have been described by which lysosomotropic compounds destabilise membranes. For instance, sunitinib and mefloquine, present among our top hits, have the ability to directly fluidise lysosomal membranes (Zhitomirsky et al., 2018). Other enhancers of antigen import, such as chlorpromazine, perphenazine, or fluphenazine have been proposed to displace lysosomal lipid hydrolases from the inner leaflet of the lysosome and to destabilise membranes by inducing changes in their lipid composition (Kornhuber et al., 2010). While the lysosomotropism of phenothiazines has been extensively studied, lysosomotropic properties of quinazolinamine-derived compounds — the most potent chemical group in the antigen import assay — has not been widely reported. Interestingly, the drug library used here includes several other lysosomotropic compounds or compounds previously shown to permeabilize lysosomes that were not active in the β-lactamase assay (e.g. norfloxacin and ciprofloxacin which destabilise lysosomes in cancer cells). Further work will be required to determine what confers selectivity of the compounds, but differences in pH, membrane composition, and proteolytic content of endolysosomal compartments are likely to influence the extent and consequences of lysosomal trapping.

Many of the clinically approved drugs demonstrate unexpected activities that have either a harmful or a beneficial effect for the patient (Pushpakom et al., 2019). These hidden effects can also be exploited for new therapeutic indications in drug repositioning approaches. Yet, predicting or identifying effects of small molecules on target cells remains challenging. Here, we took advantage of the fact that localisation and/or trafficking patterns of proteins are integral to most aspects of cellular functions. We demonstrated that comparative organellar mapping provides an effective and generic strategy for unbiased identification of biological effects of small molecules. This approach can be used to characterise changes in subcellular localisation of thousands of proteins simultaneously (without the need for antibodies or protein tagging) and to characterise on- and off-target effects in any cell type of choice (including primary cells). To our knowledge, this is also a first report of organellar mapping in cDC1s (or dendritic cells altogether), and it provides a useful resource of information on subcellular localisation and abundance of poorly characterised proteins that are not expressed in common cell lines (organellar map and full proteome composition are available at http://dc-biology.mrclmb.cam.ac.uk).

In summary, through a combination of small molecule screening and proteomics-based molecular mapping, we established a new approach for enhancing presentation of antigens sampled by DCs in the absence of strong immunogenic signals. Enhancing cross-presentation with small molecules may in the future provide therapeutic regimes for patients that do not respond to currently available treatment options.

## Supporting information

Supplemental Movei S4

## Supplemental Figures

**Figure S1.**
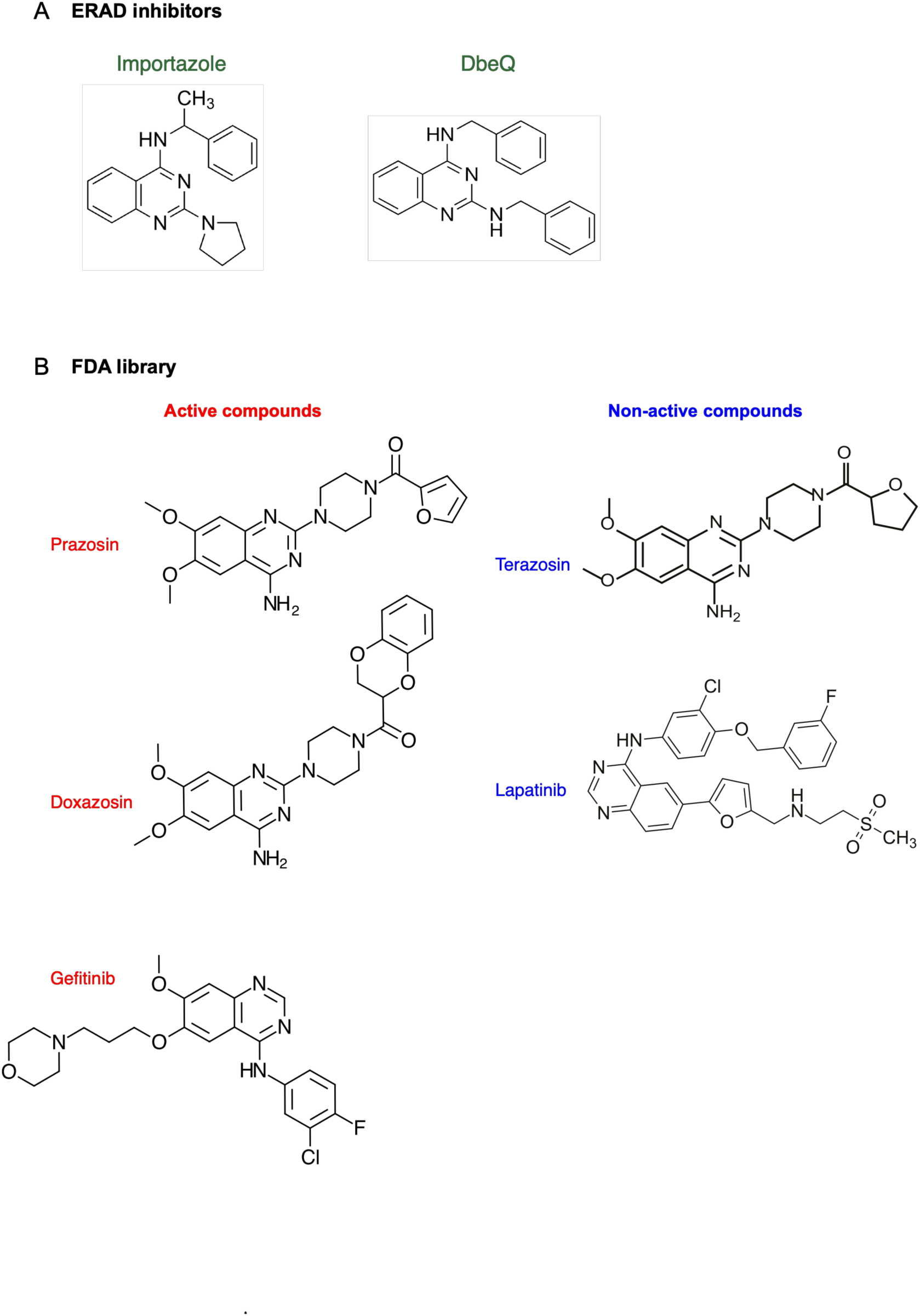
Chemical structures of the ERAD inhibitors and of selected active and non-active quinazolinamine tested in this study. *Related to Figure 1*

**Figure S2.**
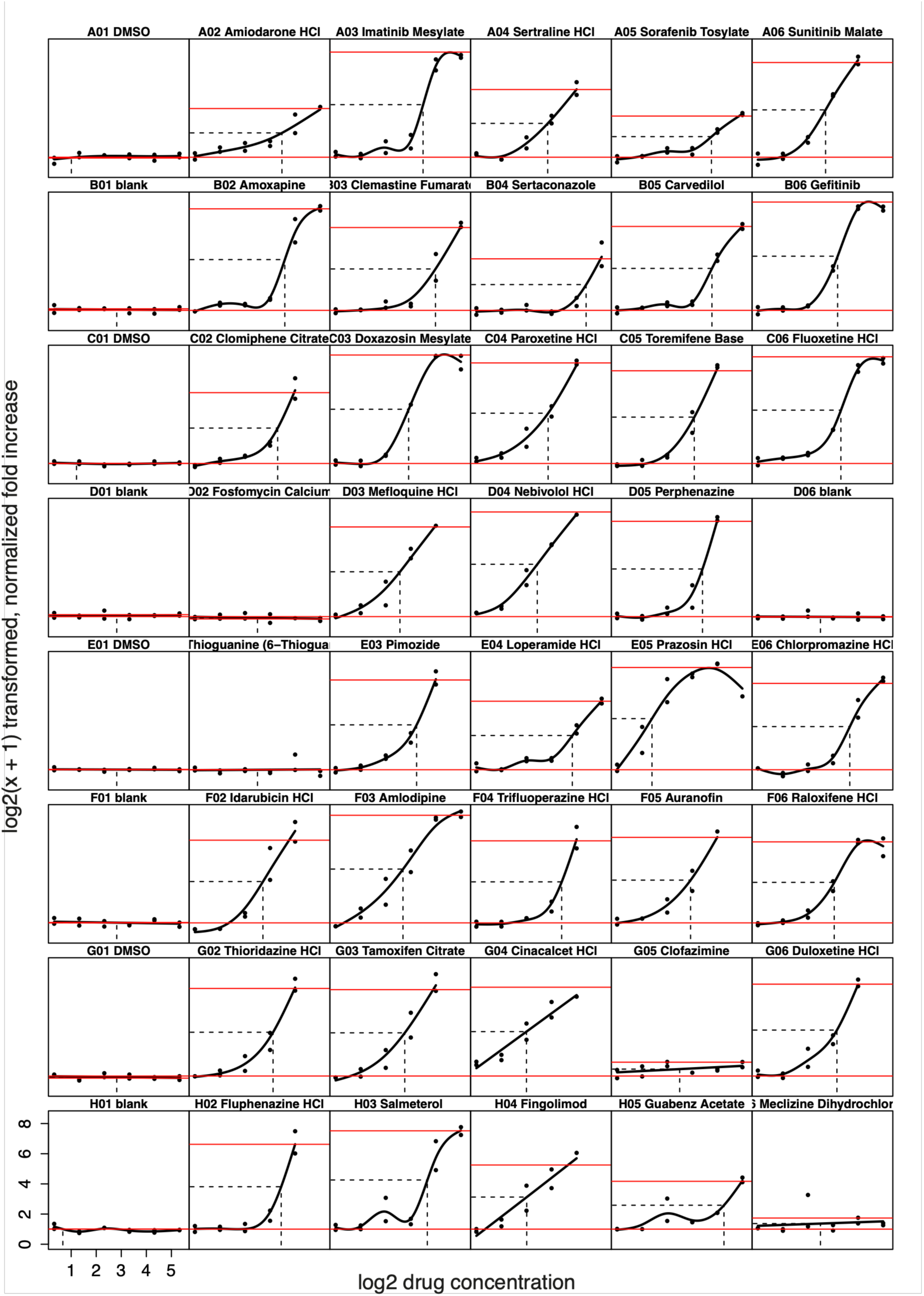
Summary of the EC50 plots from the secondary screen. *Related to Figure 1*

39 compounds were analysed using the β-lactamase assay: 37 top ranked compounds from the primary screen and two compounds with no phenotype were included (fosfomycin calcium and thioguanine). No treatment (blank) and vehicle (DMSO) controls were included on each plate. The screen was performed with five doses for each of the drugs (1.25 - 40 uM). Wells with fewer than 500 cells were excluded from analysis. The proportion of the cells with efficient β-lactamase translocation was determined and these raw phenotype measurements were normalised by dividing each value by the mean of the “DMSO” control wells from the corresponding plate. drFitSpline function from the grofit R package was used to estimate the yEC50 (50% of the max effect) values. The fold-increase and concentration values were log2(x+1) transformed for spline fitting. The concentration values were log2 transformed. The red lines indicated max and min import efficiency, the horizontal dotted line indicates the yEC50 value and the vertical line the corresponding compound concentration.

**Figure S3.**
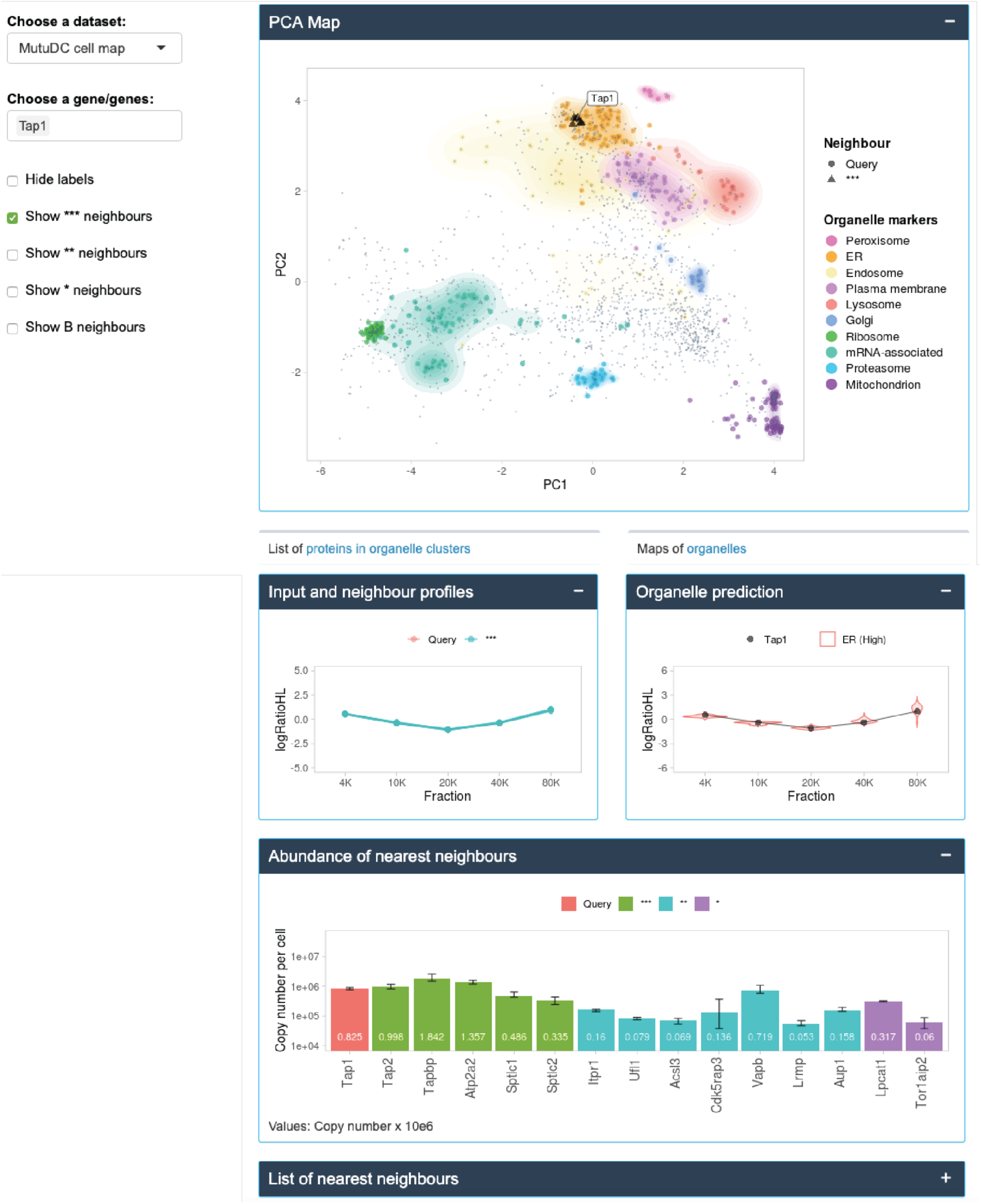
Sample output from the dc-biology.mrc-lmb.cam.ac.uk web tool for a selected gene, Tap1. *Related to Figure 1*

The maps cover over 2000 proteins expressed in MutuDCs; for each protein included in the map we identified a set of nearest neighbours and predicted the localisation. Neighbours include potential interacting partners or proteins localised to the same subcellular structure. The output of the web resource includes: representation of the query on the PCA map, profiles of the query and nearest neighbours, prediction of the subcellular localisation, absolute abundance (copy numbers and cellular concentrations), as well as a list of the nearest neighbours. In a separate section of the webtool, we provide absolute abundance for over 7000 proteins detected in MutuDCs.

**Figure S4.**
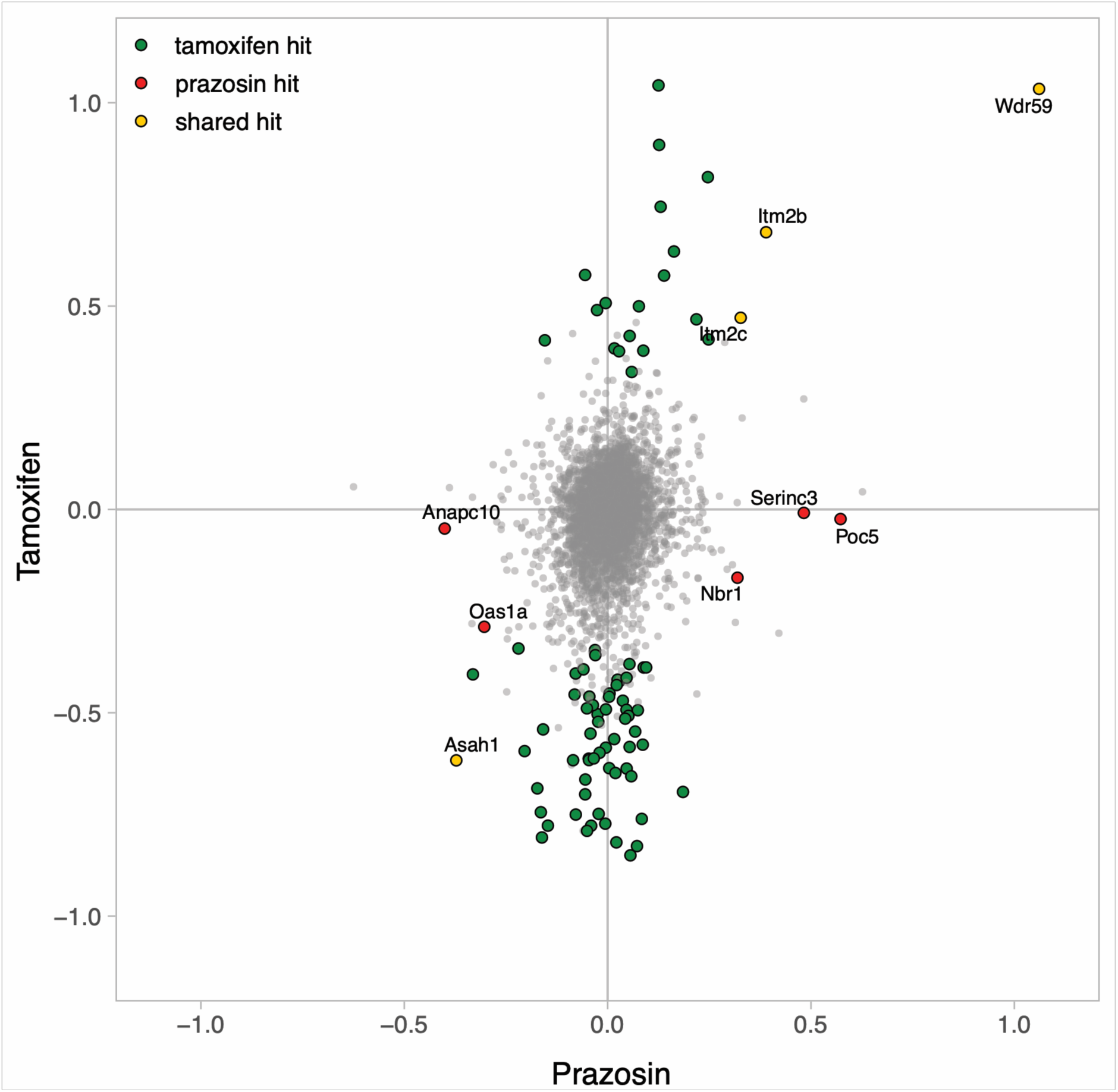
Whole cell protein expression levels in MutuDCs. *Related to Figure 4*

Protein expression levels in MutuDCs from prazosin or tamoxifen treated (4 h) cells relative to untreated controls (log 2 scale). Quantification was achieved by metabolic labelling (SILAC; averages of two replicates are shown). 5848 proteins were quantified across all four experiments. Proteins that changed significantly in abundance are highlighted in colour.

**Figure S5.**
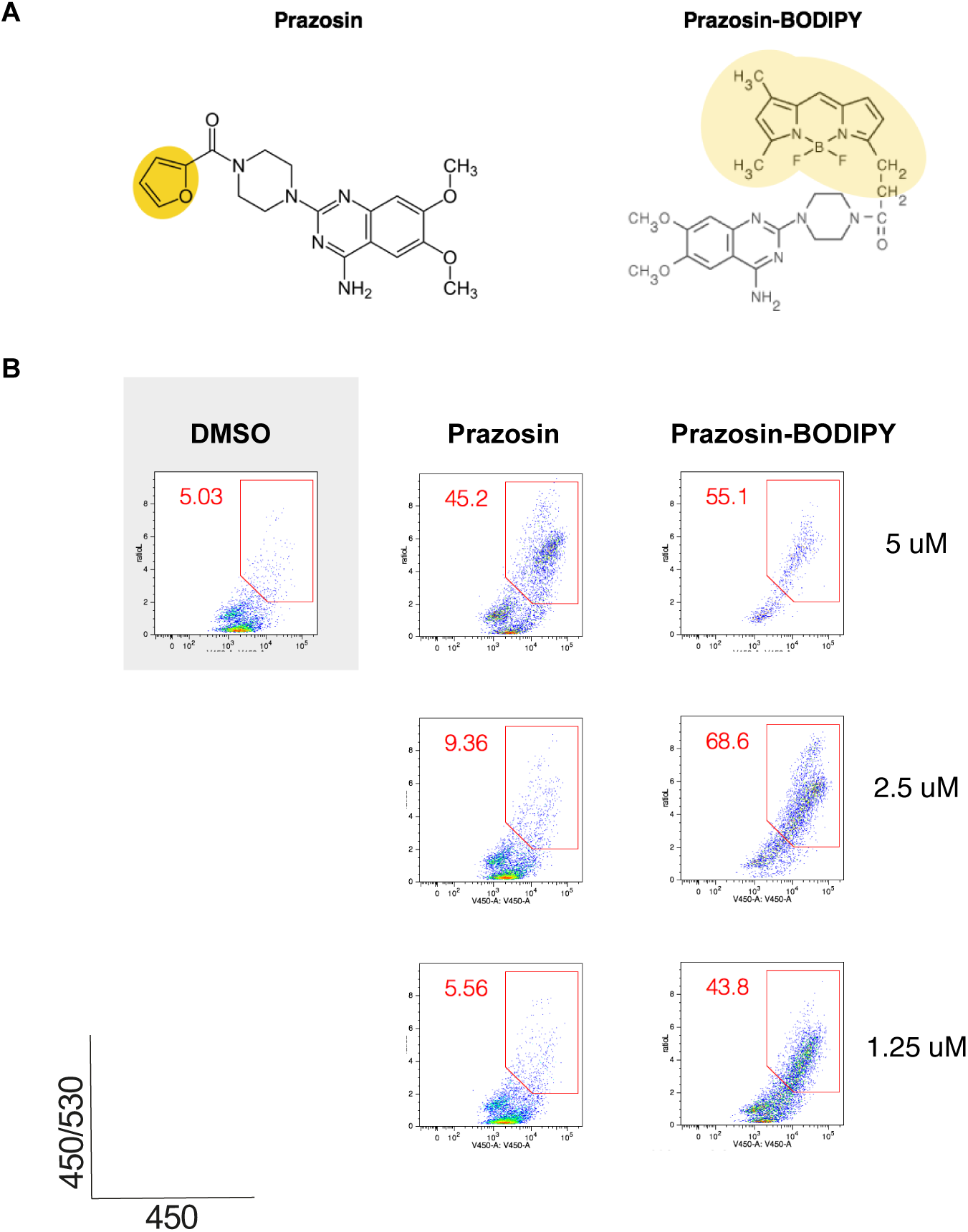
Prazosin modified with BODIPY is active in the antigen import assay. Related to Figure 5 A. Comparison of the chemical structures of prazosin and prazosin-BODIPY. B. Activity of prazosin-BODIPY in the β-lactamase assay.

## Supplemental Data

**Supplemental Data S1 - Excel file**

*Related to Figure 1*

Data from antigen import assay fοr all compounds from the FDA library

**Supplemental Data S2 – Excel file** *Related to Figures 1 and 2* Proteomic ruler in MutuDCs

MutuDC organellar map

Subcellular localisation predictions

**Supplemental Data S3 - Excel file**

*Related to Figures 3 and 4*

Organellar maps from drug-treated MutuDCs MR analysis

Whole cell and cytosol proteomics of control and drug-treated MutuDCs

**Supplemental Movie S4 - Movie**

*Related to Figure 4*

Galectin-8-YFP recruitment in control and prazosin-treated cells

## Methods

### Compounds and antibodies

For flow cytometry, the following antibodies were used: anti-CD86-PE (clone GL1, BD Pharmingen #553692), anti-CD69-PE (clone H1.2F3, BD Pharmingen #553237), anti-CD25- PerCPCy5.5 (clone PC61, BD Pharmingen #551071), anti-CD4-APC (clone RM4-5, BD Pharmingen #553051), anti-CD8α-Pacific Blue (clone 53-6.7, BD Pharmingen #558106), anti-Vα2-PeCy7 (clone B20.1, BD Pharmingen #560624), anti-CD8α-PeCy7 (clone 53-6.7, BD Pharmingen #552877), anti-Vα2-eFluor®450 (clone B20.1, eBiosciences #48-5812-82), anti-CD69-eFluor®450 (clone H1.2F3, eBiosciences #48-0691-82), anti-CD25-FITC (clone 7D4, BD Pharmingen #553072), anti-CD8a-PerCP-Cy5.5 (clone 53-6.7, eBioscience, #45-0081-82), anti-TCR vβ 5.1-PE (clone MR9-4, BD Pharmingen #553190), anti-CD4-PE-Cy7 (clone RM4-5, BD Pharmingen #552775), anti-CD19-eFluor®450 (clone 1D3, eBioscience, #48-0193), anti-CD3-eFluor®450 (clone 17A2, eBioscience, #48-0032-80), anti-CD11c-FITC (clone HL3, BD Pharmingen #553801), anti-MHC I (H-2Kb)-FITC (clone AF6-88.5.5.3, eBioscience #11-5958-80), anti-MHC II (Iab)-eFluor®450 (clone AF120.1, eBioscience #48-5320-80).

The following small molecules were used (at the indicated concentrations, unless otherwise stated in the text): DbeQ (4 μM, #SML0031), importazole (30 μM, #SML0341), PR-619 (20 μM, #SML0430), Eeyarestatin I (10 μM, #E1286), prazosin (10 μM, #P7791), prazosin (*in vivo* experiments, #1554705) tamoxifen (10 μM, #T9262), all purchased from Sigma-Aldrich; Prazosin-BODIPY (5 μM, ThermoFisher Scientific, #B7433), NMS-873 (10 μM, Selleckcheck, #S7285), SCREEN-WELL® FDA approved drug library V2 (Enzo, #BML-2843-0100).

### Animals

C57BL/6J wild type mice and C57BL/6J recombination activating gene 1 (Rag1)-deficient OT-I and OT-II TCR (Vα2, Vβ5.1) transgenic mice were obtained from Charles River Laboratories, Janvier and Centre de Distribution, Typage et Archive Animal (CDTA, Orleans, France). Mice were used between 8-12 weeks old.

All animal procedures were in accordance with the guidelines and regulations of the Institut Curie Veterinary Department and all mice used were less than six months old.

### Cell lines and cell culture

MutuDC (obtained from Hans-Acha Orbea) cells (Fuertes Marraco et al., 2012) were grown in IMDM, supplemented with 8% heat-inactivated FCS (Biowest-Biosera), 10 mM HEPES, 2 mM Glutamax, 100 IU/ml penicillin, 100 μg/ml streptomycin and 50 μM β-mercaptoethanol (all from Life Technologies).

For SILAC metabolic labelling, MutuDCs were grown in IMDM SILAC culture medium (Thermo, #88423), supplemented with 8% (V/V) dialysed fetal calf serum (PAA, #A11-107),

50 μM β-mercaptoethanol (Gibco), Pencilin and Streptomycin (Sigma), 10 mM HEPES (pH 7.4), and either: 42 mg/L 13C6,15N4-L-Arginine HCl (Silantes, #201604302) and 73 mg/L 13C6,15N2-L-Lysine HCl (Silantes, #211604302) for SILAC heavy culture medium; or 42 mg/L L-Arginine HCl and 73 mg/L L-Lysine HCl with standard isotopic constituents (Sigma, #A6969 and #L8662) for SILAC light culture medium. Cells were allowed at least seven doublings prior to experiments, to ensure complete labelling.

NIH/3T3 expressing a non-secretable form of OVA were obtained from Matthew Albert (Yatim et al., 2015), and cultured in DMEM (Life Technologies) supplemented with 10% heat-inactivated FBS (Biowest-Biosera), 0.1 mM non-essential amino acids, 1 mM sodium pyruvate, 10 mM HEPES and 50 μM β-mercaptoethanol. Necroptosis was induced by treatment with a specific drug ligand (AP20187, BB homodimerizer, Clontech).

B3Z hybridoma cells (Sanderson and Shastri, 1994) were cultured in RPMI (Life Technologies), supplemented with 10% FBS (Biowest-Biosera), 0.1 mM non-essential amino acids, 1 mM sodium pyruvate, 10 mM HEPES and 50 μM β-mercaptoethanol, 10 mM HEPES.

B16-OVA cells (Falo et al., 1995) were cultured in RPMI, supplemented with 10% heat-inactivated FCS (Biowest-Biosera), 2 mM Glutamax, 100 IU/ml penicillin and 100 μg/ml streptomycin (all from Life Technologies) and selected with G418 2 mg/ml (Life Technnologies) and hygromycin B 60 μg/ml (Gibco).

MC38-OVA (Gilfillan et al., 2008) cells were grown in DMEM, supplemented with 10% heat-inactivated FCS (Biowest-Biosera), 2 mM Glutamax, 100 IU/ml penicillin and 100 μg/ml streptomycin (all from Life Technologies).

OT-I and OT-II T cells were isolated using EasySep™ Mouse Naïve CD8+ and CD4+ T Cell Isolation Kits respectively (Stemcell, #19858 and #19765) and cultured in the same media as the B3Z cells.

All cell lines testes as mycoplasma-negative by PCR.

### Antigen import assay and library screen

MutuDCs were seeded at 150,000 cells/well in U-bottom 96-well plates and incubated with 10 mg/ml β-lactamase (Sigma, #P0389) for 3 h at 37°C. The cells were then washed and incubated with small molecules at indicated concentrations for 2 h at 37°C. CCF4 loading was performed as described1 for 45-60 min at RT. To increase the sensitivity of the assay, the plates were then incubated for 16 h at RT21 in CO2 independent media supplemented with 8% FCS, and 2 mM Glutamax. Immediately before the flow cytometry analysis, the cells were stained with Fixable Viability Dye eFluor® 780 (eBioscience) diluted 1:2500 in PBS. Proportion of the live cells with a high ratio of blue to green (V450/V530) fluorescence was used as a measure of the efficiency of antigen import into the cytosol.

Primary screen of the FDA library. Enzo FDA-approved drug library was screened in the course of three independent experiments. Each 96-well plate contained three media-only,

DMSO only, 4 μΜ DbeQ (enhancement control), and 10 μM PR-619 (inhibition control) wells to control for data reproducibility between the plates. The screen was performed once and 37 top ranked compounds were selected for validation.

Validation screen. The secondary screen was performed at six concentrations (1.25 – 40 μM) for each compound, in two biological repeats. Media-only and vehicle (DMSO) controls were included on each plate. Wells with less than 500 cells were excluded from analysis. The raw phenotype measurements (percent of cells with a high ratio of blue-to-green fluorescence) were normalised by dividing each value by the mean of media-only control wells from the corresponding plate. The EC50 values were estimated using a drFitSpline function from the grofit R package. (log2(x+1) transformed values were used for spline fitting). Note that for some drugs the max effect might not have been reached at the maximum concentration tested, which might result in underestimation of the EC50 values).

### Chemical class and target assignment

The information about chemical classes and candidate targets was downloaded from DrugBank database (Law et al., 2014). The enrichment of chemical classes and targets in active vs non-active compound groups was calculated using Fisher’s test (R studio). Only the primary target was selected for each drug for the enrichment analysis.

### Antigen presentation assays

#### Cross-presentation assay

1×105 MutuDC were seeded in round bottom 96-well plates and incubated with different concentrations of soluble grade VII OVA (Sigma Aldrich #A7641) or endotoxin-free OVA (Hyglos #300036). Minimal peptide OVA_257-264_ was used as a control for the capacity of DCs to activate T cells. As indicated, MutuDCs were either incubated with OVA for 45 min, followed by a 3.5 h incubation with small molecules or incubated with OVA and small molecules continuously for 5h. Next, DCs were washed three times with 0.1% (vol/vol) PBS/BSA, fixed with 0.008% (vol/vol) glutaraldehyde for 3 min at 4°C, washed twice with 0.2 M glycine and twice with the T growth media. 1×105 B3Z hybridoma cells were added per well. After 16 h, the cells were lysed in a buffer containing 9 mM MgCl_2_, 0.125% NP40 (Nonidet® P40 substitute, Santa Cruz #sc-29102) 1.7 mM chlorophenol red-β-D-galactopyranoside (CPRG, Roche #10884308001). CPRG conversion by β-galactosidase was measured by optical density at 590 nm.

#### OTI and OTII activation assays assay

1×104 DCs per well were seeded in round bottom 96-well plates and incubated for 5h with different concentrations of grade VII OVA (Sigma Aldrich #A7641), endotoxin-free OVA (Hyglos #300036), or control minimal peptides (OVA_257-264_ and OVA_323-339_). Where indicated, prazosin was added at 10 μM or Poly(I:C) at 5 μg/ml. After 5h, DCs were washed three times with PBS containing 0.1% (vol/vol) BSA and co-cultured with purified OT-I CD8^+^ or OT-II CD4+ T for 16h. For monitoring T cell activation, T cells were stained for CD69 and CD25 and analysed by flow cytometry.

#### Cell-associated antigens

1×105 MutuDC were seeded in round bottom 96-well plates with 3T3-OVA cells at various 3T3-OVA:MutuDC ratios (1:2, 1:4, 1:8, 1:16, 1:32). The co-cultures were incubated at 37°C in the presence of prazosin (10 μM) or DMSO (1:1000). After 5h, the co-cultures were washed, fixed and co-incubated with B3Z hybridomas or washed and co-cultured for 16h with 1×105 purified OT-I or OT-II T cells. B3Z and OTI/II T cell activation was monitored as described above.

### Cell uptake assay

NIH/3T3 were stained with the PKH-26 membrane dye (Sigma Aldrich, #PKH26-GL) following the manufacturer’s instructions. 1×105 MutuDC were plated in 96 round bottom-well plates together with PKH-26+ 3T3s at different 3T3:MutuDC ratios (1:2, 1:4, 1:8, 1:16, 1:32). The co-cultures were incubated in the presence of prazosin (10 μM) or DMSO for 2h or 5h at 37°C, 5% CO2, or left on ice for 5h. Cells were then stained with anti-CD11c-APC (clone HL3, BD Pharmingen #550261) and violet live/dead Dye (ThermoFisher #L34955) and fixed to prevent further uptake.

Percentage of PKH-26+ MutuDCs (CD11c+ cells) was determined. Phagocytic index was calculated by subtracting the percentage of PKH-26+ cells in CD11c+ gate obtained at 4°C from the percentage of this subset measured at 37°C after 2h or 5h of incubation.

### DC activation

To assess DC activation, 1×105 MutuDC were seeded in round bottom 96-well plates and incubated for 5h with endotoxin-free OVA (Hyglos #300036) or grade VII OVA (Sigma Aldrich #A7641-250MG), in presence or absence of DMSO or prazosin (10 μM). After 5h, cells were washed twice with medium, cultured for additional 15h, and finally stained for CD86.

### Live microscopy

Images were acquired on a VisiTech iSIM swept field confocal super resolution system coupled to a Nikon Ti2 inverted microscope stand equipped with a 100x/1.49 NA SR Apo TIRF objective lens. Fluorophores were excited simultaneously with 488 nm and 561 nm laser light and imaged with two Hamamatsu ORCA-Flash4.0 V3 CMOS cameras via an image splitter (filter: ZT561rdc from Chroma Technology). MutuDCs were seeded the day before the experiment. The imaging was performed at 37°C with 5% CO_2_.

### Tumour growth experiments

MC38-GFP-OVA. WT mice were injected subcutaneously with 2x106 OVA-expressing MC38 cells 100 μl of cold-sterile 1× PBS. The experiment was then conducted as described previously. Tumour growth was measured three times a week and volume was calculated as (height × width2)/2 (where width is the shorter measurement). When tumour size reached 1000 mm3, the mice were euthanised.

B16-OVA. WT mice were injected subcutaneously with 2.5x105 OVA-expressing B16 cells in 100 μl of cold-sterile PBS. When tumours became visible, usually within a week, mice were randomly assigned to different treatment groups. Injections of prazosin (0.5 mg/mouse) and/or anti-PD1 antibody (200 μg/mouse) were then performed three times per week, starting the day of tumour appearance. Vehicle (cold water and/or PBS) was injected into control mice. Tumour growth was measured three times a week and volume was calculated as above. When tumour size reached 1000 mm3, mice were euthanized. To control for toxic effects of prazosin, we performed a pilot experiment in which mice were treated for a period of one month with: 0.5 g prazosin in 1 ml, administered IP, 3x a week (total of 13 injections, total dose: 7.5 g prazosin per mouse); no adverse effects were observed.

The mean growth rate curves were estimated using loess function in R. The statistical significance analysis was performed in Prism using ANOVA with FDR Benjamini-Hochberg correction.

### Whole cell proteomics

MutuDCs were grown in SILAC light or SILAC heavy culture medium, in 15 cm dishes, to 70-90% confluency. SILAC heavy labelled cells were treated with tamoxifen or prazosin (20 μM) for ~4 h at 37 °C; SILAC light labelled cells were treated with DMSO (vehicle) only. Cells were incubated for 3h 45 min at 37 °C, and harvested. Cell pellets were lysed in SDS buffer (2.5% (w/V) SDS, 50 mM Tris-HCl, pH=8.0), and incubated at 90°C for 10 min. To shear genomic DNA, lysates were passed through a QIAashredder (Qiagen). Lysates were then processed for analysis by mass spectrometry as described below. For the repeat experiment, the SILAC labelling of control and treated cells was swapped. Protein concentrations were estimated by BCA assay. Equal amounts of control and treated samples (i.e. SILAC light and heavy, or vice versa) were pooled, and acetone precipitated as described (Itzhak et al., 2016). Samples were subjected to tryptic digest using the FASP method (Wiśniewski et al., 2009). Peptides were fractionated into six fractions using strong cation exchange (Kulak et al., 2014) (SCX), prior to mass spectrometric analysis.

### Proteomic analysis of cytosol

MutuDCs were cultured in SILAC light or SILAC heavy growth medium, in 10 cm dishes, to 70-90% confluency. SILAC light cells were treated with tamoxifen, prazosin (both at 10 μM), or vehicle (DMSO) for ~4 h at 37 °C; SILAC heavy labelled cells were left untreated, and served as internal reference. Treatments were performed in quadruplicate (two pairs of replicates prepared on two different days). Cells were harvested and resuspended in STE buffer (250 mM sucrose, 0.5 mM MgCl_2_, 0.2 mM EGTA, 25 mM Tris-HCl, pH = 7.5 at 4°C). Aliquots of SILAC heavy labelled cells were mixed with proportional aliquots of the tamoxifen-, prazosin-or DMSO-treated SILAC light cells. Cells were lysed mechanically in a Dounce homogenizer (tight pestle, 40 strokes, on ice). Lysates were centrifuged at 2,000 x g for 10 min at 4°C, to pellet cell debris und nuclei. Post nuclear supernatants were centrifuged at 135,000 x g for 45 min at 4°C to pellet organelles and microsomes. Supernatants were the cytosolic fraction. Protein concentrations were estimated by BCA assay; aliquots were acetone precipitated and subjected to in-solution digest and stage-tip peptide cleanup as previously described (Itzhak et al., 2016), prior to mass spectrometric analysis.

### Dynamic organellar maps

Organellar maps were prepared essentially as described (Itzhak et al., 2016), with minor modifications to the protocol. Briefly, MutuDCs were cultured in SILAC light or SILAC heavy growth medium, in 15 cm dishes, to 70-90% confluency. SILAC light cells were treated with tamoxifen or prazosin (10 μM), or vehicle (DMSO), for 4 h; SILAC heavy labelled cells were treated with vehicle (DMSO), and served as reference. Two dishes were used for each treatment (SILAC light cells), and four dishes to generate the SILAC heavy reference. Unlike in (Itzhak et al., 2016), the same reference was used for treated and control maps. Cells were harvested (with the drugs or DMSO added to the PBS (-) cell detachment buffer), chilled on ice, lysed mechanically in STE buffer (250 mM sucrose, 0.5 mM MgCl_2_, 0.2 mM EGTA, 25 mM Tris-HCl, pH = 7.5 at 4°C), with a Dounce homogenizer, and centrifuged at 1,000 x g for 10 min to pellet nuclei and cell debris. Post-nuclear supernatants of SILAC light labelled cells were then subjected to a series of differential centrifugation steps (4,000 x g for 10 min; 10,000 x g for 15 min; 20,000 x g for 20 min; 40,000 x g for 20 min; 80,000 x g for 30 min). Post nuclear supernatant from SILAC heavy cells was centrifuged once at 80,000 x g for 30 min to obtain the reference fraction. All pellets were resuspended in SDS buffer (2.5% (w/V) SDS, 50 mM Tris-HCl, pH=8.0), and heated to 90 °C for 3 min. Protein concentrations were estimated by BCA assay. Equal amounts of SILAC heavy reference fraction were mixed with each SILAC light subfraction, acetone precipitated and subjected to in-solution digest and stage-tip peptide cleanup as described (Itzhak et al., 2018), prior to mass spectrometric analysis.

Fractionations were prepared in duplicate, on two different days (six maps total – two controls, two from cells treated with tamoxifen, and two from cells treated with prazosin).

### Mass spectrometry and data processing

Mass spectrometric analysis was performed as described (Itzhak et al., 2016), using a Thermo EASY-nLC 1000 HPLC coupled to a Q Exactive HF Hybrid Quadrupole-Orbitrap (Thermo Fisher Scientific, Germany). HPLC gradient lengths varied for the different experiments. For analysis of whole proteomes, each of the SCX peptide fractions was analysed with a 240 min gradient (24 h per sample in total). For the analysis of cytosol and fractions from the organellar maps, each sample was analysed with a single 150 min gradient. Raw files were processed with MaxQuant software Version 1.6 (Cox and Mann, 2008), using the murine reference proteome (Swiss-Prot canonical and isoform data) database downloaded from UniProt (www.uniprot.org).

### Bioinformatic analysis of the proteomic data

Protein groups identified through MaxQuant analysis were filtered to remove reverse hits, proteins identified with modified peptides only, as well as common contaminants. Further processing depended on the individual experiment:

#### Copy number estimates of proteins expressed in MutuDC

To estimate absolute protein abundance in MutuDCs, the SILAC datasets used for full proteome analysis of drug-treated cells were used (see below). Each of the four dataset combined control cells and drug treated cells. From each dataset, the protein intensities from the control cells were selected, to obtain four replicate full proteomes. Intensities within each replicate were summed, and all replicates were linearly normalized to the same summed intensity. Next, only proteins detected in at least two replicates were retained (7427 in total). Copy number estimates were calculated using the Proteomic Ruler (Wiśniewski et al., 2014), as implemented in Perseus software (.5, (Tyanova et al., 2016), and described in (Itzhak et al., 2018). Protein intensities were scaled to molecular mass.

#### Drug-induced changes in whole cell proteomes

For analysis of drug-induced changes in whole cell proteomes, only proteins with at least three SILAC quantification events in each of the four experiments (2 x control vs tamoxifen, 2 x control vs prazosin) were retained (5848 proteins). SILAC ratios were linearly normalized to a column median of 1 in each experiment, logarithmised, and analysed with the ‘Significance A’ tool in Perseus software (Tyanova et al., 2016). Proteins that changed significantly in both replicate experiments with one drug (FDR=0.05 within each replicate, Benjamini-Hochberg correction), with a consistent direction of change, were considered as hits for this drug. Proteins that changed significantly across both replicates and both drug treatments, with a consistent direction of change in all four measurements, were considered as hits common to both drugs.

#### Drug-induced changes in cytosol

For analysis of drug-induced changes in cytosol, only proteins with at least three SILAC quantification events in each of the four replicates were retained (2129 proteins). SILAC ratios were linearly normalized to a column median of 1 in each experiment, and logarithmised. For each protein, the average ratio SILAC light/SILAC heavy from the four replicates was calculated for each condition, and average control (DMSO) ratios were then subtracted from average treatment (tamoxifen or prazosin) ratios. Thus, for every protein, the average change in cytosolic levels caused by either tamoxifen or prazosin relative to DMSO was obtained.

The log ratios from the whole cell proteome and cytosol analyses were plotted against each other for each treatment (including only proteins detected in both). To compare distribution of lysosomal proteins with distribution of all detected proteins the Kolmogorov–Smirnov test was performed.

#### Organellar maps

Generation of organellar maps and outlier testing followed the principles described in detail (Itzhak et al., 2018; 2016), with some modifications to accommodate a comparison across three conditions. Only proteins with high quality SILAC ratios in all 30 subfractions, i.e. across all six maps, were retained (1857 proteins). (High quality SILAC ratios are those calculated from three or more quantification events. In addition, ratios calculated from only two quantification events are also included in the high quality set if the corresponding MaxQuant ratio variability is below 30%). Each map consisted of a set of five SILAC ratios for each protein, mirroring its distribution across the differential centrifugation fractions. SILAC ratios were inverted, and divided by the sum of all five ratios across the map. This yielded for each protein a ‘per map’ normalised profile (summing to 1). For the MutuDC control map shown in Figure 2C/3B, all proteins passing the high quality filter in both replicates were included (2121 proteins). To visualize the map the prcomp function in R was used, with the following parameters: (center = TRUE, scale. = TRUE). Organellar marker proteins were initially chosen from our previously published set, and augmented as described (Itzhak et al., 2016).

#### Subcellular localisation predictions in MutuDC

Organellar maps from the two control map replicates (0-1 normalised, Supplemental Data S2) were annotated with 559 markers for 12 organellar compartments, by cross-matching our previously derived set of human marker proteins (Itzhak et al., 2016). Support vector machines (implemented in Perseus software,.6, (Tyanova et al., 2016)) were trained to predict organellar association as described ((Itzhak et al., 2016)), with an overall recall of 93% and a median F1 score of 0.88 across all compartments (Supplemental Data S2).

#### Drug induced protein movements

To identify proteins that moved significantly and robustly, our previously reported MR (movement and reproducibility (Itzhak et al., 2016)) analysis was applied, with minor modifications. Unlike in our previous study, here only one reference fraction was used to generate the control and treatment maps. This reference came from cells treated with DMSO only. A different normalization was therefore required, to allow the outlier test to detect changes in membrane association as well as organellar localization shifts. SILAC ratios were first normalised within each fraction to a column median of 1. Next, for each protein, SILAC ratios were inverted, and weighted with fraction yields (determined by BCA assay (Itzhak et al, 2016). Within each map all data were then summed. This reflected overall amount of protein detected in each map (prep yield). The smallest prep yield was set to one, and correction factors for the other five maps were calculated relative to this value. All data within a map were then globally normalised through division by the prep yield correction factor. The result were six maps in which the sum of all data points is equal. Next, for each protein the ten data points from the two tamoxifen replicates and the ten corresponding data points from control replicates were divided by the sum of all of these ratios. The same was repeated for the ten data points from the two prazosin replicates, using the same ten control data points. This procedure results in an additional “within treatment” normalisation of the maps. Next, for each protein, the treatment profiles were subtracted from the corresponding control profiles, to yield ‘delta’ profiles. For every protein, four delta profiles, with five data points each (two sets from tamoxifen and two sets from prazosin treatment) were obtained. Delta profiles from treatment replicates were combined into one profile (ten data points) and analysed with the multivariate outlier test in Perseus software (Perseus 1.6, 101 iterations, quantile = n*0.75)

(Itzhak et al., 2016). Movement (M) scores were calculated as the negative log(10) of the FDR corrected p values (Benjamini-Hochberg method). For example, an M score of four identifies significantly moving proteins with an FDR of 0.01%. The reproducibility (R) score was calculated as the Pearson correlation of the two five-data point delta profiles within treatment replicates. A significance cut-off corresponding to a p-value of 0.05 (R=0.8) was chosen. Since the R-score represents an additional filter, orthogonal to the M-score, further multiple hypothesis correction of the p-value was not required. Each protein with significant M (>4) and R (>0.8) scores was considered as shifting significantly. Thus, for every protein two sets of M and R scores were obtained, reflecting shifts caused by tamoxifen or prazosin treatment. Each treatment produced a partially overlapping list of shifting proteins.

## Acknowledgements

We would like to thank Hans Acha-Orbea for the MutuDCs, Matthew Albert for the NIH/3T3 cells expressing a non-secretable form of OVA, Felix Randow for the Galectin-3-YFP construct, and Jon Howe for help with microscopy. S.A. received funding from: Institute Curie; Institut National de la Santé et de la Recherche Médicale; Centre National de la Recherche Scientifique; la Ligue Contre le Cancer (Equipe labellisée Ligue, EL2014.LNCC/SA); Association de Recherche Contre le Cancer (ARC); the ERC (2013-AdG N° 340046 DCBIOX), INCA PLBIO13-057; ANR-11-LABX-0043 and ANR-10-IDEX-0001-02 PSL; ANR-16-CE15001801 and ANR-16-CE18002003. P.K. was supported by EMBO (ALTF 467-2012), the

Wellcome Trust (101578/Z/13/Z) and Medical Research Council (U105178805). A.A. was supported by EMBO (ALTF 883-2011). DNI was funded by the Louis-Jeantet Foundation, and the Max Planck Society for the Advancement of Science; GHHB was funded by the German Research Foundation (DFG/Gottfried Wilhelm Leibniz Prize), and the Max Planck Society for the Advancement of Science. DNI and GHHB also wish to thank Matthias Mann for his support.

## Author contributions

P.K performed, analysed, designed, and supervised experiments; M.G. performed and analysed part of the antigen presentation assays, set up and carried out experiments with cell-associated antigens. D.N.I. carried out the proteomic experiments, and assisted with data analysis. L.J. performed tumour experiments and assisted with all animal work. M.G., and A.A assisted with tumour experiments. J.G.M. assisted in assay development and experimental design. D.N.E. assisted with design of the small molecule screening assays. G.H.H.B. designed the proteomics experiments, performed the data analysis, and supervised the proteomics work. P.K. and S.A. conceived and supervised the study; and P.K., G.H.H.B and S.A. wrote the manuscript.

